# Aging disrupts cumulus–oocyte NAD⁺ homeostasis

**DOI:** 10.64898/2026.07.22.739977

**Authors:** Bettina P. Mihalas, Denise Lin, Sonia Bustamante, Russell Pickford, Emily R. Frost, Ananya Vuyyuru, Derek Y. Wong, Michael J. Bertoldo, Lindsay E. Wu, Robert B. Gilchrist

## Abstract

The age-related decline in oocyte nicotinamide adenine dinucleotide (NAD⁺) is associated with reduced developmental potential and female infertility. Despite the extraordinary longevity of the female germline, the mechanism for maintaining oocyte NAD⁺ remains unresolved. Here, we used stable isotope tracing to identify a new mechanism for shared, intercellular NAD^⁺^ biosynthesis, whereby somatic–germline metabolic coupling between the oocyte and its surrounding cumulus cells is critical to maintain oocyte NAD^⁺^ homeostasis. We show that this coupling deteriorates with reproductive aging, identifying altered NAD⁺ metabolism in cumulus cells from mice and women of advancing reproductive age. This cumulus–oocyte metabolic coupling of NAD⁺ biosynthesis contributes to protection against the age-related increases in oocyte reactive oxygen species (ROS). In intact complexes, restoring NAD⁺ through supplementation with the precursor nicotinamide mononucleotide (NMN) increased glutathione, reduced ROS and improved mitochondrial membrane potential in oocytes from aged mice and in oocytes exposed to oxidative insult. Importantly, the ability of NMN to resolve elevated ROS depends on the presence of cumulus cells. Together, this new model of somatic-germline metabolic coupling of NAD^⁺^ biosynthesis places an age-related deterioration in cumulus cell-mediated metabolic support as a key driver of impaired oocyte NAD^⁺^ levels and redox dysregulation with aging.

## Introduction

Female infertility represents the first manifestation of aging in mammals, preceding the decline in all other physiological systems^1,2^. This is primarily driven by declining oocyte quality, which is a core feature of reproductive aging and the major cause of age-related female infertility^3,4^. In humans from the mid-thirties onward, declining oocyte quality increases the time to conception and the risk of miscarriage and stillbirth^5–7^, and advanced maternal age has also been associated with adverse outcomes in offspring^8–14^. This decline is of enormous clinical significance - despite up to 64% of assisted reproductive technology (ART) cycles being undertaken by women aged 35 years or older, current ARTs are unable to reverse the age-related decline in oocyte quality^3,4^.

An age-associated decrease in nicotinamide adenine dinucleotide (NAD⁺) has been reported across a spectrum of cell and tissue types in multiple organisms including humans and rodents^15–17^. Declining NAD⁺ levels in mouse^18–20^ and human oocytes^21^ during reproductive aging have emerged as an important contributor to declining oocyte quality^18,20,22^. NAD⁺ is a redox coenzyme that is central to cellular energy metabolism and mitochondrial function^23,24^. NAD⁺ acts as an electron acceptor during the oxidation of glucose, fatty acids and tricarboxylic acid cycle intermediates. Its reduced form, NADH, subsequently donates these electrons to the mitochondrial electron transport chain to support oxidative phosphorylation and adenosine triphosphate (ATP) production^23^. The phosphorylated redox couple, nicotinamide adenine dinucleotide phosphate (NADP⁺/NADPH), provides reducing power for antioxidant defense, including the regeneration of reduced glutathione and thioredoxin, and fuels anabolic processes such as *de novo* lipogenesis^25,26^. NAD⁺ also acts as a substrate for NAD⁺-dependent enzymes, including sirtuins, poly(ADP-ribose) polymerases (PARPs) and CD38, which contribute to metabolic regulation, redox homeostasis, epigenetic control, DNA repair and immune regulation^23^.

In many tissues, neighboring cells exhibit metabolic cooperation, exemplified by neuron–astrocyte coupling^27^, Sertoli–germ cell coupling^28^, and photoreceptor–Müller glia–retinal pigment epithelial cell metabolic exchange^29^. The ovarian follicle represents a highly specialized form of this intercellular cooperation, most notably within the cumulus–oocyte complex (COC), where the oocyte and surrounding cumulus cells function as a syncytium. Within the COC, cumulus cells extend transzonal projections (TZPs) across the zona pellucida forming gap junctions at the oocyte membrane^30^, to facilitate the transfer of nucleotides, amino acids, and sugars^31,32^. The COC presents an extreme example of metabolic coupling, in which the oocyte is dependent on surrounding somatic cells to accumulate the cytoplasmic stores that are essential for oocyte growth, meiotic regulation, and early embryonic development^31^. The oocyte alone is unable to carry out central metabolic pathways, including glycolysis, and relies on cumulus cells to take up glucose, metabolize it to pyruvate, and supply pyruvate to the oocyte to support oxidative phosphorylation^31,32^. Cumulus cells have also been proposed to directly transport NADPH into the oocyte^33,34^. Taken together, it is clear that cumulus–oocyte communication within the COC is central to oocyte developmental competence^35^.

Restoration of NAD⁺ levels through treatment with its precursors nicotinamide mononucleotide (NMN) and nicotinamide riboside^18,20,22^, along with the inhibition of the NAD⁺-consuming enzyme CD38^36,37^, have been shown to improve oocyte quality and fertility in reproductively aged mice, and protect against chemotherapy-induced infertility^38^. Ten days of intraperitoneal NMN injection improved oocyte mitochondrial activity and ATP levels, reduced ROS, and improved spindle structure and chromosome alignment, reduced aneuploidy, and improved embryo development^20^, and four weeks of oral NMN supplementation in aged mice improved oocyte yield, spindle structure, oocyte size, embryo quality and breeding performance^18^. In the aging ovary, declining NAD^⁺^ levels are linked to increased CD38-mediated NAD^⁺^ consumption, associated largely with infiltration of immune cells into the ovarian stroma^36,37^. While this may explain declining NAD^⁺^ across the entire ovary, the cause of declining NAD⁺ within the oocyte specifically, the rate-limiting cell for fertility, remains unresolved.

During reproductive aging, cumulus–oocyte communication deteriorates. Aging is associated with slowed cumulus cell expansion^39^, fewer TZPs and reduced gap-junction coupling^40^, and altered cumulus cell molecular signatures^41–44^. The functional significance of these age-related changes is supported by recent work showing that aged mouse oocytes reconstructed within a young somatic environment displayed improved maturation, blastocyst formation and live-birth outcomes^45^. Together, these findings suggest that age-related deterioration of cumulus–oocyte communication may contribute to the decline in oocyte developmental competence.

In this study, we investigate the role of cumulus cells in regulating oocyte NAD⁺ homeostasis during reproductive aging. We present a model of shared intercellular NAD⁺ biosynthesis in which somatic–germline metabolic coupling supports the conversion of NMN into oocyte NAD⁺. This cumulus–oocyte interaction is critical for assimilation of exogenous NMN into oocyte NAD^⁺^, and for the ability of NMN to resolve elevated oocyte reactive oxygen species (ROS). Finally, we show that NAD⁺ metabolism is altered in mouse and human cumulus cells during reproductive aging. Together, these findings identify age-related deterioration of cumulus cell-mediated metabolic support as a cause of declining oocyte NAD⁺ homeostasis during reproductive aging, placing the cumulus cells as a therapeutic target to restore oocyte quality in aging.

## Results

### Reproductive aging alters NAD⁺ and adenine metabolite dynamics within the COC

In this study, we sought to determine whether altered cumulus–oocyte metabolic coupling during reproductive aging influences oocyte NAD⁺ metabolism, hypothesizing that accelerated metabolic aging of cumulus cells contributes to impaired oocyte NAD⁺ homeostasis. To test this, we first quantified NAD⁺ metabolites in dissociated COCs from reproductively young and old mice (Fig. 1). We utilized the SwissTacAusb strain, as this model parallels the age-related decline in oocyte yield and increased aneuploidy documented in humans^46^. We measured NAD⁺ salvage pathway metabolites, which are essential for NAD^⁺^ generation and developmental competence in the oocyte^18,47^ (Fig. 1a). Intact COCs from young (5–7 weeks) or reproductively aged (12–14 months) mice were mechanically separated to obtain denuded oocytes and their corresponding cumulus cells, which were subjected to targeted quantitative metabolomics. This revealed an age-related decline in multiple NAD⁺ metabolites in oocytes (Fig. 1b), including nicotinamide (NAM), NADP^⁺^ and NADPH, as well as a trend toward decreased NAD^⁺^. In this context, a notably increased NADPH/NADP⁺ ratio in old oocytes likely reflects disproportionate NADP⁺ depletion rather than enhanced antioxidant capacity. When we examined matched cumulus cells, we were surprised to find that the somatic compartment of the COC also displayed reduced NAD⁺ metabolites with reproductive aging, including NADP⁺ and NAD⁺ (Fig. 1c).

**Fig. 1:**
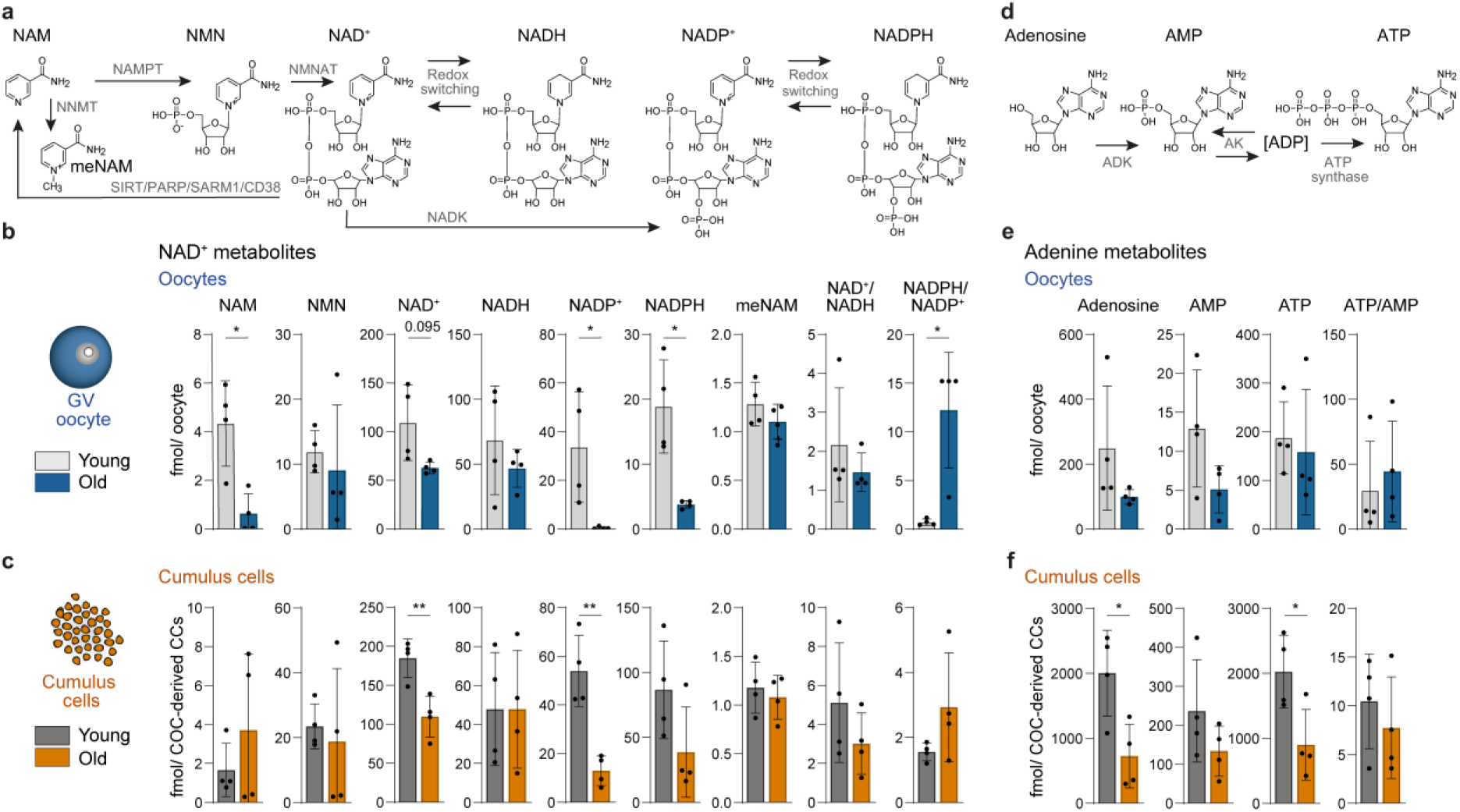
Reproductive aging alters NAD⁺ and adenine metabolites in mouse cumulus–oocyte complexes. **a,** Schematic of NAD⁺ metabolism. **b,c,** NAD⁺-related metabolites measured by targeted LC–MS/MS in **b,** germinal vesicle (GV)-stage oocytes and **c,** matched cumulus cells isolated from cumulus–oocyte complexes (COCs) from reproductively young (5–7 wk) and old (12–14 mo) mice. **d,** Schematic of adenine nucleotide metabolism. **e,f,** Adenine metabolites measured by targeted LC–MS/MS in **e,** GV oocytes and **f,** matched cumulus cells from the same experimental groups. Metabolite abundance is shown as fmol per oocyte for oocyte samples and fmol per COC-derived cumulus cells (CCs) for cumulus cell samples. Bars show mean ± s.d.; individual dots represent independent biological replicates (n = 4). Each biological replicate comprised pooled oocytes or matched cumulus cells from 20–30 COCs collected from a minimum of three mice. For normally distributed data, statistical significance was determined using two-sided unpaired Welch’s t-tests. Where one or both groups failed normality testing, two-sided Mann–Whitney tests were used. Asterisks indicate statistical significance: \**P* < 0.05 and \*\**P* < 0.01. Exact *P* values are shown for comparisons with *0.05 < P* < 0.1; unlabeled comparisons had *P* ≥ 0.1.

Given the interlinked nature of NAD⁺ and adenine metabolism, whereby ATP contributes the adenylyl moiety to NAD⁺ biosynthesis^48^, and NAD⁺ and NADH support ATP generation^23^, we also examined adenine metabolites in these same samples (Fig. 1d). Although no detectable age-related differences in adenine metabolites were observed in oocytes (Fig. 1e), cumulus cell adenosine and ATP levels more than halved with age (Fig. 1f).

We further examined NAD⁺ and adenine metabolite dynamics during meiotic maturation across reproductive age by comparing immature germinal vesicle (GV)-stage samples with mature metaphase II (MII)-stage samples (Extended Data Fig. 1). In young oocytes, meiotic maturation to the MII stage corresponded with decreased NAD⁺, NADP⁺ and NADPH, an increased NADPH/NADP⁺ ratio, and a trend toward decreased NADH (Extended Data Fig. 1a), likely reflecting consumption of these metabolites during the energy-demanding process of meiosis^49^. Notably, these meiotic maturation-associated changes were not similarly observed in reproductively aged oocytes, where only a trend toward decreased NMN was detected, potentially reflecting reduced NAD⁺ turnover in aged oocytes. Cumulus cells also displayed modest changes during meiotic maturation. In young cumulus cells, there was a trend toward increased NAM and decreased NADPH during maturation, whereas in aged cumulus cells, NMN and the NADPH/NADP⁺ ratio decreased (Extended Data Fig. 1b).

For adenine metabolites, there were no detectable changes in oocytes across meiotic maturation (Extended Data Fig. 1c); however, adenosine increased in cumulus cells from both reproductively young and old COCs, indicating enhanced purine/adenine nucleotide turnover during meiotic maturation (Extended Data Fig. 1d).

To test whether these age-related changes were conserved in human cumulus cells, NAD⁺ and adenine metabolites were examined in leftover cumulus cells donated by patients of different ages undergoing routine intracytoplasmic sperm injection (ICSI) treatment, during which the cumulus cell layer is routinely removed prior to sperm injection, and discarded. This analysis highlighted an inverse correlation between patient age and levels of NAM, meNAM and adenosine (Fig. 2a and Fig. 2b). A similar reduction in NAD^⁺^ levels with age was not observed, however, the strong age-dependent reduction in nicotinamide and 1-methyl-nicotinamide but not NAD^⁺^ likely reflects the limits of sample preservation for clinical samples (Fig. 2a). NAD^⁺^ can rapidly degrade into nicotinamide and then 1-methyl-nicotinamide if not subject to rapid freezing and/or metabolite extraction, however, these samples were obtained from a working IVF clinic in which leftover cumulus cells were collected hours after they had been removed from patient oocytes, with highly variable times of sample collection. Together, these findings suggest that age-related changes in cumulus cell NAD⁺-linked metabolism are partially conserved between mice and humans, extending our mouse findings into a clinically relevant human context.

**Fig. 2:**
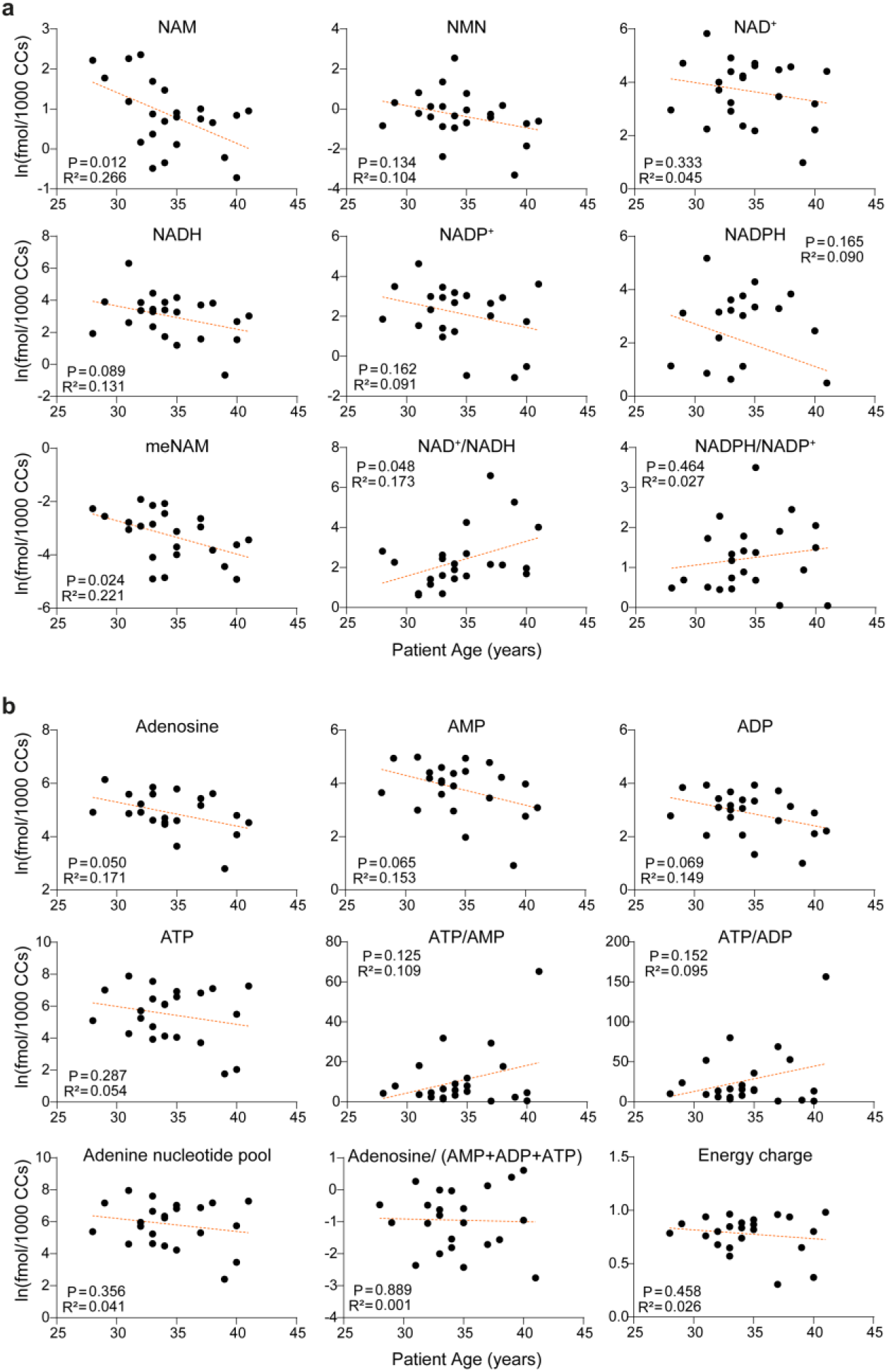
NAD⁺ and adenine metabolites are altered in human cumulus cells with increasing reproductive age. **a,b,** Targeted LC–MS/MS quantification of **a,** NAD⁺-related metabolites and **b,** adenine metabolites in peri-ovulatory cumulus cells donated by patients undergoing intracytoplasmic sperm injection treatment. Samples were obtained from 23 patients. Each dot represents cumulus cells from one patient. Metabolite abundance was normalized to live-cell number and is shown as fmol per 1,000 live cumulus cells (CCs). Associations between patient age and log-transformed metabolite abundance were assessed by simple linear regression. Dashed orange lines show simple linear regression of log-transformed metabolite abundance against patient age. P values and R² values for each metabolite are shown in each graph.

These data demonstrate that age-related metabolic changes in the NAD⁺ metabolome are not restricted to the oocyte, supporting the idea that accelerated metabolic aging of cumulus cells may contribute to impaired oocyte NAD⁺ metabolism.

### Cumulus–oocyte communication is required for oocyte NAD⁺ homeostasis

Having identified a decline in cumulus cell NAD⁺ metabolites with reproductive age, and given that aging is associated with fewer TZPs and reduced gap-junction coupling in the COC^40^, we sought to test whether cumulus cells contribute to the maintenance of oocyte NAD⁺ homeostasis. To test this, cumulus–oocyte communication within the COC was disrupted using two orthogonal approaches. Transport from cumulus cell transzonal projections into the oocyte was blocked through broad chemical inhibition of cumulus–oocyte gap junctions using the small molecule inhibitor carbenoxolone (CBX). In parallel, cumulus–oocyte communication was blocked by the mechanical removal of cumulus cells from the oocyte, resulting in denuded oocytes, which do not receive cumulus cell contact and support. COCs were maintained under 3-isobutyl-1-methylxanthine (IBMX)-mediated meiotic arrest for 17 h in the presence or absence of NMN, allowing cumulus–oocyte communication to be examined under conditions in which TZP-mediated connectivity remains high. Oocytes and matched cumulus cells were then isolated for targeted metabolomics (Fig. 3a).

**Fig. 3:**
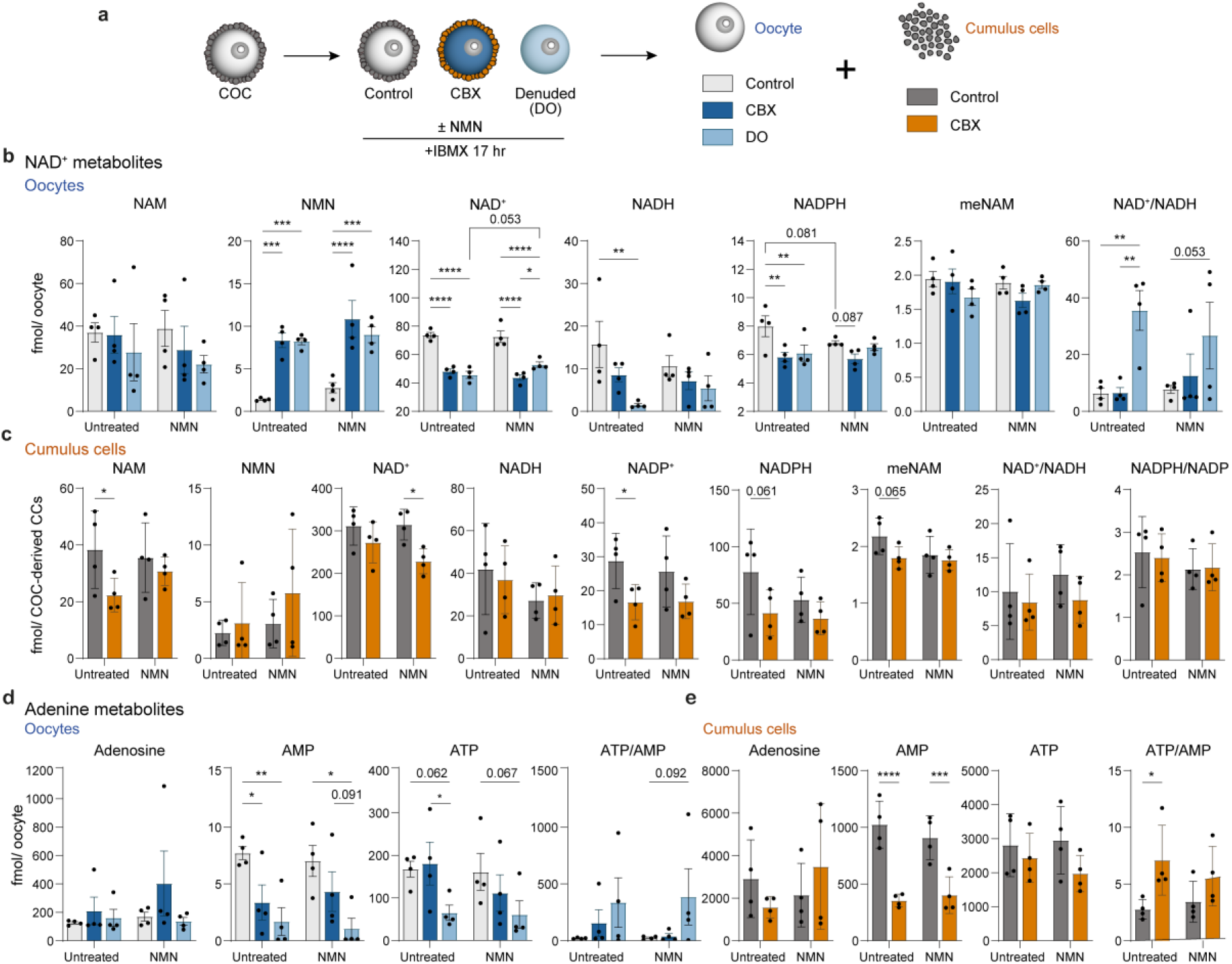
Cumulus–oocyte communication regulates oocyte NAD⁺ and adenine metabolism. **a,** Experimental overview. Germinal vesicle (GV) cumulus–oocyte complexes (COCs) from reproductively young mice (5–7 wk) were cultured intact as controls, or treated with carbenoxolone (CBX) to inhibit gap-junction communication, or mechanically denuded to remove cumulus cells (denuded oocytes; DOs), in the presence or absence of nicotinamide mononucleotide (NMN) for 17 h under 3-isobutyl-1-methylxanthine (IBMX)-mediated meiotic arrest. Oocytes and cumulus cells were then isolated for targeted LC–MS/MS. **b,** NAD⁺-related metabolites measured in oocytes from control and CBX-treated COCs and DOs. **c,** NAD⁺-related metabolites measured in matched cumulus cells from control and CBX-treated COCs. **d,** Adenine metabolites measured in oocytes from control and CBX-treated COCs and DOs. **e,** Adenine metabolites measured in matched cumulus cells from control and CBX-treated COCs. Metabolite abundance is shown as fmol per oocyte for oocyte samples and fmol per COC-derived cumulus cells (CCs) for cumulus cell samples. Bars show mean ± s.d.; individual dots represent independent biological replicates (n = 4). Each biological replicate comprised pooled oocytes or matched cumulus cells from 20–30 COCs collected from a minimum of 12 mice. Statistical significance was determined using ordinary two-way ANOVA. For oocyte analyses, COC communication condition and NMN treatment were used as factors; for cumulus cell analyses, CBX treatment and NMN treatment were used as factors. Both analyses were followed by Fisher’s LSD test for selected row- and column-wise pairwise comparisons. Asterisks indicate statistically significant selected comparisons: \**P* < 0.05, \*\**P* < 0.01, \*\*\**P* < 0.001 and \*\*\*\**P* < 0.0001. Exact *P* values are shown for selected comparisons with *0.05 < P* < 0.1.

Disruption of cumulus–oocyte communication led to pronounced changes in both oocyte and cumulus cell NAD⁺ metabolism. In oocytes, both CBX treatment and mechanical denudation reduced NAD⁺ (Fig. 3b). Strikingly, this depletion in NAD^⁺^ levels following the disruption of cumulus–oocyte communication led to a sharp accumulation in the levels of the immediate NAD^⁺^ precursor NMN. These data suggest that the final step of oocyte NAD^⁺^ biosynthesis, the conversion of NMN to NAD^⁺^, is critically dependent on interactions with cumulus cells. Notably, exogenous NMN did not restore NAD⁺ levels in CBX-treated or denuded oocytes, further reinforcing that cumulus cell communication is critical to oocyte NAD⁺ homeostasis.

Matched cumulus cells from CBX-treated COCs also displayed altered NAD⁺ metabolism, with reduced NAM and NADP⁺, trends toward reduced NADPH and meNAM, and reduced NAD⁺ in NMN-supplemented CBX-treated cumulus cells (Fig. 3c).

For adenine metabolites, disruption of cumulus–oocyte communication reduced oocyte AMP in denuded oocytes and CBX-treated oocytes, while ATP abundance was not significantly altered relative to control oocytes (Fig. 3d). Exogenous NMN did not restore these adenine metabolite changes. Similarly, CBX caused a marked reduction in cumulus cell AMP, whereas ATP was maintained (Fig. 3e), resulting in an increased ATP/AMP ratio in CBX-treated cumulus cells consistent with impaired adenine nucleotide cycling^50^.

To further examine whether the cumulus-enclosed state supports oocyte metabolism, we compared oocytes isolated from intact COCs (Fig. 1) to ‘naked’ oocytes which were naturally recovered without surrounding cumulus cells, rather than having had the cumulus cell layer deliberately removed through mechanical denuding, as in Fig. 3 (Extended Data Fig. 2a). Naked oocytes are not uncommon in clinical IVF cycles. Compared to intact COCs collected from the same cycle, naked oocytes displayed reduced abundance of most NAD⁺ metabolites, including NAM, NAD⁺, NADH, NADPH and meNAM, with a trend toward decreased NADP⁺ (Extended Data Fig. 2b). Despite reduced NADPH, the NADPH/NADP⁺ ratio was increased in naked oocytes, potentially reflecting a disproportionate loss of NADP⁺ rather than enhanced reductive capacity. AMP was also reduced in naked oocytes, while ATP abundance was maintained (Extended Data Fig. 2c). Together, these changes in oocyte NAD^⁺^ metabolism following disruption of cumulus–oocyte communication support a model of intercellular NAD^⁺^ homeostasis.

Given that disruption of cumulus–oocyte communication results in NMN accumulation and NAD⁺ depletion (Fig. 3b), we next tested the idea that these changes were the direct result of impaired conversion of NMN into NAD^⁺^. We measured the assimilation of NMN into NAD^⁺^ by tracing the incorporation of stable isotope-labeled NMN, in which the NAM moiety is labeled with four deuterium atoms (d₄-NMN). Following the assimilation of labeled NMN into NAD⁺, cellular redox cycling can exchange one deuterium at the NAM redox site, such that the incorporation of the label is detected predominantly as d₃-NAD⁺ (Fig. 4a)^51,52^. Consistent with this, treatment of intact COCs with d₄-NMN resulted in the formation of d₃-NAD⁺ as the predominant NAD⁺ isotopologue and displaced unlabeled endogenous NAD^⁺^ (Extended Data Fig. 3). Strikingly, NAD^⁺^ labeling was vastly reduced in both CBX-treated and denuded oocytes. As in previous results (Fig. 3), treatment with CBX or mechanical denudation led to the accumulation of NMN, in both its labeled and endogenous unlabeled forms. This shows that the reduction in NAD^⁺^ labeling during the disruption of cumulus–oocyte communication is not due to impaired oocyte NMN uptake, as NMN uptake can occur independently of cumulus cells; however, the conversion of this NMN into NAD^⁺^ is critically dependent on cumulus cell interactions (Fig. 4b,c). Due to this labeling strategy in which the loss of one deuterium label indicates prior assimilation into NAD^⁺^, the absence of changes in endogenous or labeled d₃-NAM and d₃-NADP⁺ suggests that labeled metabolites can still enter the NAD⁺ salvage pathway, but are not efficiently converted into NAD⁺ in the oocyte.

**Fig. 4:**
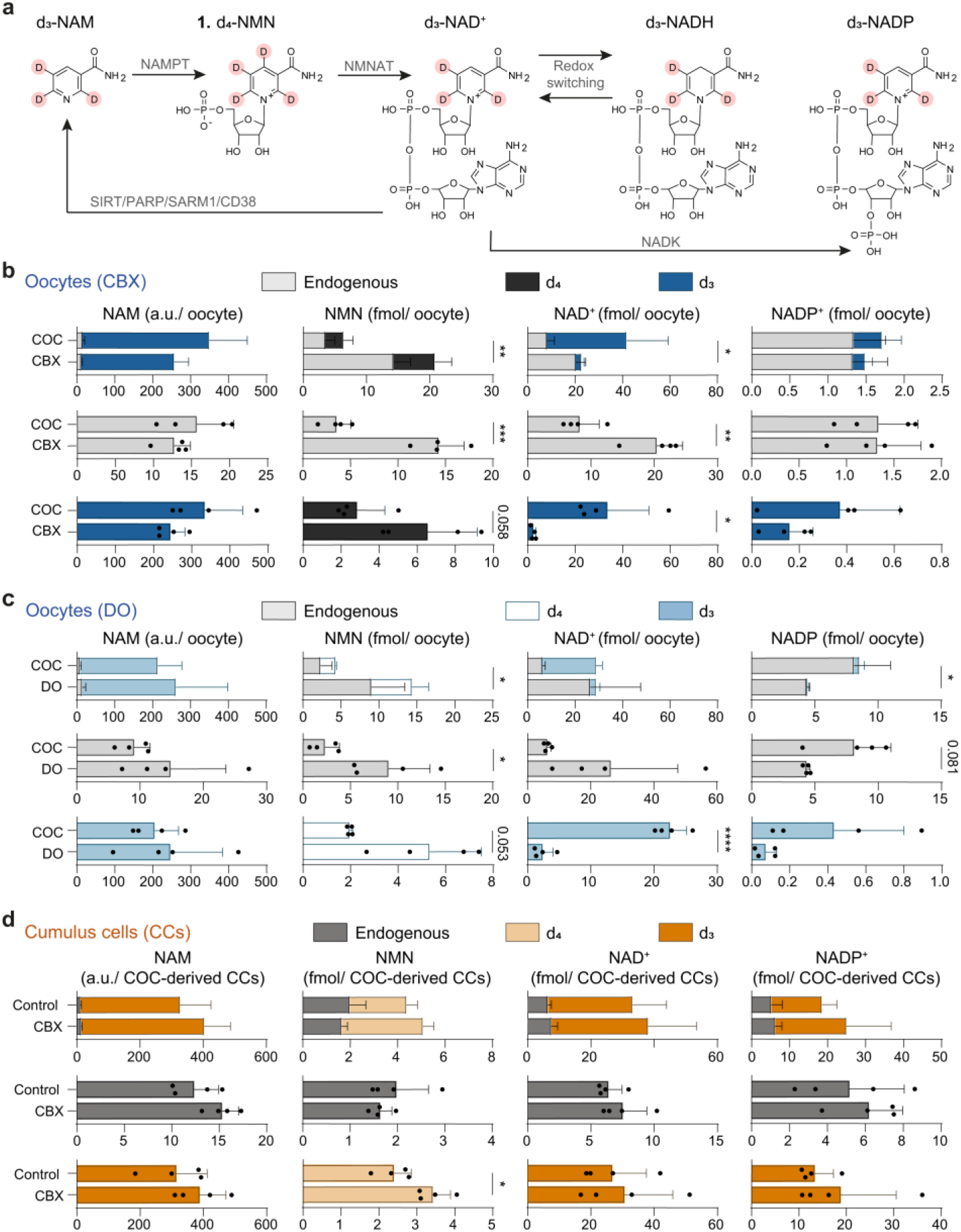
Cumulus–oocyte communication is required for exogenous NMN to support NAD⁺ synthesis in oocytes. **a,** Schematic of deuterium-labeled NMN tracing through the NAD⁺ metabolic pathway. Germinal vesicle (GV)-stage cumulus–oocyte complexes (COCs) from reproductively young mice (5–7 wk) were cultured intact as controls, treated with carbenoxolone (CBX) to inhibit gap-junction communication, or mechanically denuded to remove cumulus cells (denuded oocytes; DOs), in the presence of deuterium-labeled NMN (d₄-NMN) for 17 h under isobutyl-1-methylxanthine (IBMX)-mediated meiotic arrest. **b,c,** Endogenous and deuterium-labeled NAD⁺-related isotopologues (d₄ and d₃) measured by targeted LC–MS/MS in oocytes following **b,** CBX treatment or **c,** mechanical denudation. **d,** Endogenous and deuterium-labeled NAD⁺-related isotopologues (d₄ and d₃) measured by targeted LC–MS/MS in matched cumulus cells from COCs treated ± CBX. Metabolite abundance is shown as fmol per oocyte for oocyte samples and fmol per COC-derived cumulus cells (CCs) for cumulus cell samples except for nicotinamide (NAM), which is shown as relative abundance (a.u.). Bars show mean ± s.d.; individual dots represent independent biological replicates (n = 4). Each biological replicate comprised pooled oocytes or matched cumulus cells from 40–50 COCs collected from a minimum of six mice. For normally distributed data, statistical significance was determined using two-sided unpaired Welch’s t-tests. Where one or both groups failed normality testing, two-sided Mann–Whitney tests were used. Asterisks indicate statistical significance: \**P* < 0.05, \*\**P* < 0.01, \*\*\**P* < 0.001 and \*\*\*\**P* < 0.0001. Exact *P* values are shown for comparisons with *0.05 < P < 0.1*; unlabeled comparisons had *P* ≥ 0.1.

In contrast to the oocyte, matched cumulus cells from CBX-treated COCs did not show the same loss of labeled NAD⁺ synthesis (Fig. 4d). Instead, cumulus cells maintained d₃-NAD⁺ and d₃-NADP⁺ abundance following CBX treatment, suggesting that they remain competent to take up and metabolize exogenous NMN when gap-junction communication is inhibited. Similar to oocytes, however, d₄-NMN also accumulated in CBX-treated cumulus cells. Of note, 94–96% of the detected NAM pool was d₃-NAM across oocytes and cumulus cells, indicating that the majority of NAM was derived from metabolized exogenous NMN. This is consistent with rapid NMN uptake and active NAD⁺ salvage pathway turnover within the COC. Preservation of d₃-NAD⁺ synthesis in cumulus cells following gap-junction inhibition, together with the greater capacity of cumulus cells to synthesize d₃-NADP⁺ relative to oocytes, highlights distinct NAD⁺ metabolite dynamics between the two cell types.

### NMN mitigates age-related oxidative stress in cumulus-enclosed oocytes

Elevated ROS is a major metabolic feature of aged oocytes^5^ and is linked to oxidative stress-associated defects in chromosome alignment^49,53^, increased aneuploidy^54,55^ and impaired embryonic development^56^. Given the central role of NAD⁺ metabolism in redox homeostasis, and our finding that cumulus–oocyte communication supports oocyte NAD⁺ synthesis, we next examined whether age-related disruption of cumulus cell NAD⁺ metabolism compromises the capacity of the COC to support oocyte redox balance.

We chose to use an *in vitro* maturation (IVM) system, allowing us to isolate cumulus–oocyte communication from the broader ovarian environment while simultaneously assessing the consequences for mature MII oocyte quality and redox homeostasis. COCs from reproductively young and old mice underwent IVM in the presence or absence of NMN, before isolation of MII oocytes and cumulus cells for targeted metabolomics (Fig. 5a). In MII oocytes, reproductive aging was associated with reduced NMN and NAD⁺, with a trend toward reduced NADPH under NMN-treated conditions (Fig. 5b). Aging was also associated with a reduced NAD⁺/NADH ratio, consistent with a shift in oocyte redox state. Remarkably, NMN supplementation increased NAD⁺ in old oocytes, but not young oocytes, indicating that age-related decline in oocyte NAD⁺ metabolism can be partially improved during IVM in cumulus-enclosed oocytes.

**Fig. 5:**
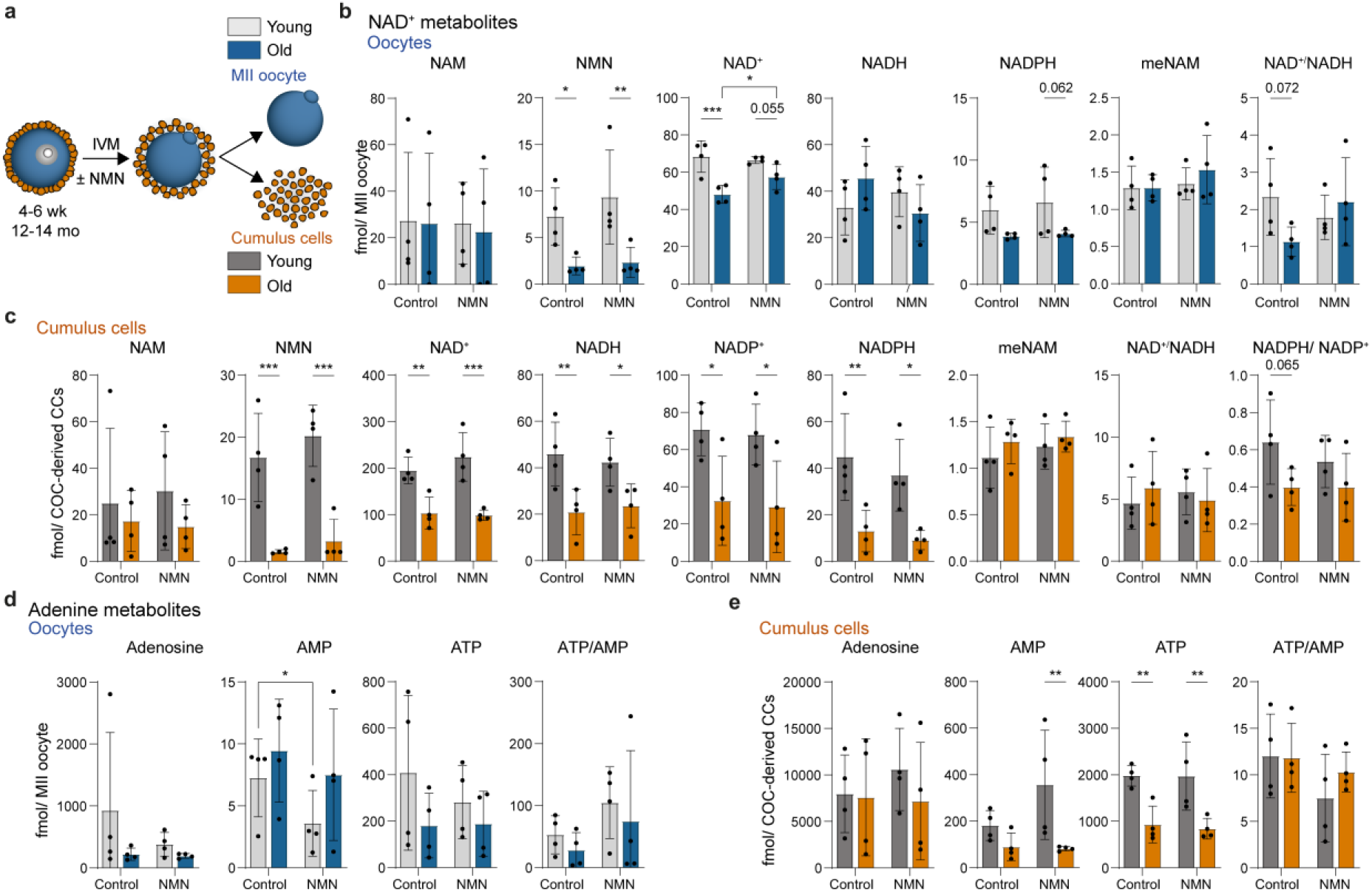
NMN partly mitigates age-related changes in NAD⁺ and adenine nucleotide metabolism in maturing cumulus-enclosed oocytes. **a,** Experimental overview. Cumulus–oocyte complexes (COCs) from reproductively young (5–7 wk) and old (12–14 mo) mice underwent in vitro maturation (IVM) in the presence or absence of nicotinamide mononucleotide (NMN), before isolation of metaphase II (MII)-stage oocytes and matched cumulus cells for targeted LC–MS/MS. **b,c,** NAD⁺-related metabolites measured in **b,** MII oocytes and **c,** matched cumulus cells. **d,e,** Adenine metabolites measured in the same **d,** MII oocytes and **e,** matched cumulus cells. Metabolite abundance is shown as fmol per MII oocyte for oocyte samples and fmol per COC-derived cumulus cells (CCs) for cumulus cell samples. Bars show mean ± s.d.; individual dots represent independent biological replicates (n = 4). Each biological replicate comprised pooled oocytes or matched cumulus cells from 20–30 COCs collected from a minimum of three mice. Statistical significance was determined by using repeated-measures two-way ANOVA, with age and NMN treatment as factors and matching by biological replicate across NMN treatment within each age group, followed by row- and column-wise pairwise comparisons using Fisher’s LSD test. Asterisks indicate statistically significant selected comparisons: \**P* < 0.05, \*\**P* < 0.01 and \*\*\**P* < 0.001. Exact *P* values are shown for selected comparisons with *0.05 < P* < 0.1.

Cumulus cells displayed larger and broader age-related changes than oocytes in NAD⁺ metabolism, including reduced NMN, NAD⁺, NADH, NADP⁺ and NADPH, as well as a trend toward a reduced NADPH/NADP⁺ ratio (Fig. 5c). These age-related changes were not restored by NMN supplementation.

For adenine metabolites, MII oocytes showed no detectable age-related differences (Fig. 5d). However, AMP was significantly reduced in NMN-treated young MII oocytes, potentially reflecting altered adenine nucleotide balance associated with the ATP-dependent conversion of NMN to NAD⁺^48^. In cumulus cells, aging was associated with reduced ATP abundance, whereas AMP was reduced only in NMN-treated cumulus cells from old mice compared with the corresponding young group (Fig. 5e). This, together with the wholesale loss of NAD⁺ metabolites in old cumulus cells (Fig. 5c), reveals significant impairment of cumulus cell metabolism in older mice by the end of oocyte maturation.

The partial restoration of NAD⁺ abundance in old oocytes following NMN treatment (Fig. 5b) was also reflected by improved oocyte redox homeostasis in MII-stage oocytes (Fig. 6). In our model, aging increased polar body extrusion rates, which may be consistent with an age-related weakening of spindle assembly checkpoint–mediated meiotic arrest (Fig. 6b)^57^. The expected age-related increase in intra-oocyte ROS (CM-H₂DCFDA; Fig. 6c) and decrease in glutathione (monochlorobimane; Fig. 6d), were both ameliorated with NMN supplementation during IVM. In addition, the age-associated decline in polarized mitochondria (J-aggregates) and mitochondrial membrane potential (J-aggregate/J-monomer) were significantly improved by the addition of NMN (Fig. 6e). However, NMN supplementation decreased both JC-1 monomer and J-aggregate fluorescence in young oocytes without altering the J-aggregate/J-monomer ratio.

**Fig. 6:**
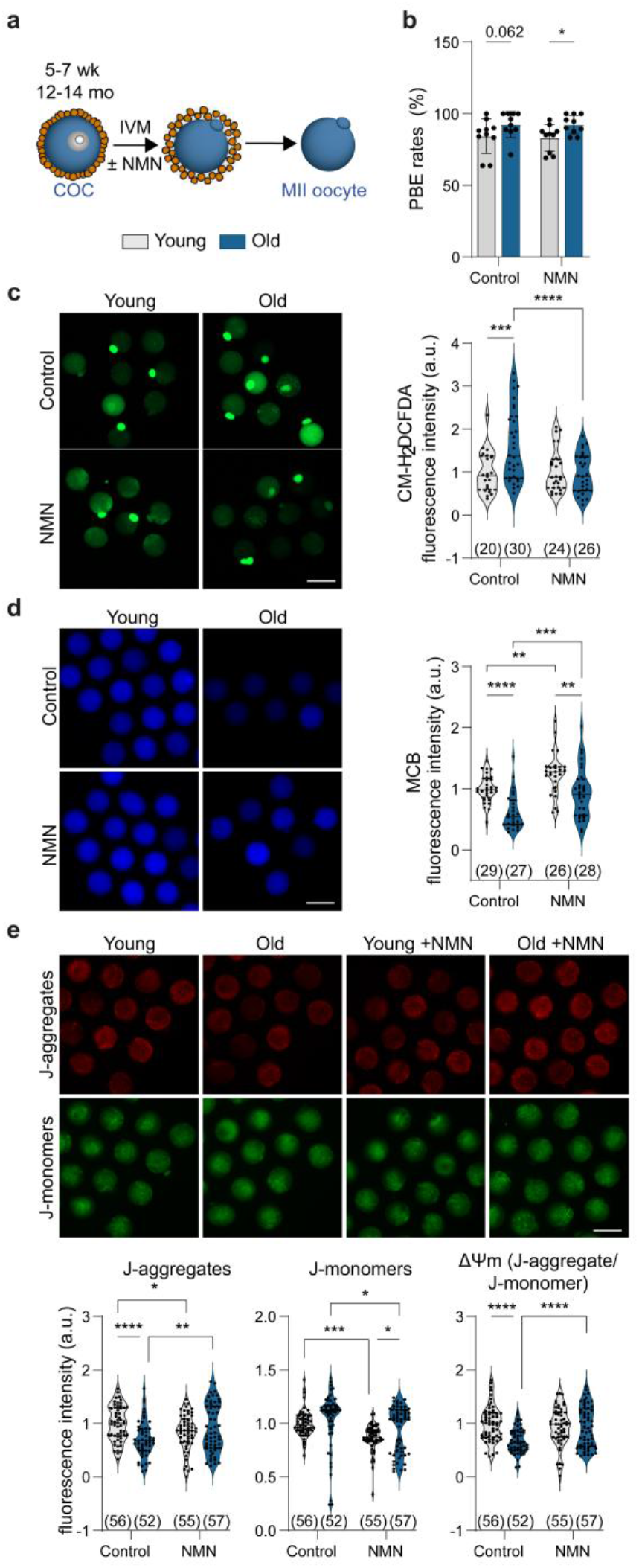
NMN mitigates age-related oxidative stress in cumulus-enclosed oocytes. **a,** Experimental overview. Cumulus–oocyte complexes (COCs) from reproductively young (5–7 wk) and old (12–14 mo) mice underwent in vitro maturation (IVM) in the presence or absence of nicotinamide mononucleotide (NMN). **b,** Polar body extrusion (PBE) rates following IVM. **c,d,** Representative images and quantification of **c,** CM-H₂DCFDA fluorescence intensity (indicative of reactive oxygen species) and **d,** monochlorobimane (MCB) fluorescence intensity (indicative of reduced glutathione) in metaphase II (MII) oocytes. **e,** Representative images and quantification of JC-1 red aggregate fluorescence, green monomer fluorescence and red:green fluorescence ratio in MII oocytes, indicative of mitochondrial membrane potential. Scale bars, 100 µm. Fluorescence intensity is shown in arbitrary units (a.u.) for all fluorescence analyses. Bars show mean ± s.d. For PBE analyses, each dot represents an independent biological replicate, with each replicate containing a minimum of 10 oocytes. For fluorescence analyses, each dot represents an individual oocyte from a minimum of three biological replicates, with each replicate containing a minimum of four imaged oocytes. Oocyte numbers are annotated on the x-axis within each graph. For all experiments, oocytes were collected from a minimum of three mice. Statistical significance was determined using ordinary two-way ANOVA, with age and NMN treatment as factors, followed by Fisher’s LSD test for selected row- and column-wise pairwise comparisons. Asterisks indicate statistically significant selected comparisons: \**P* < 0.05, \*\**P* < 0.01, \*\*\**P* < 0.001 and \*\*\*\**P* < 0.0001. Exact *P* values are shown for selected comparisons with *0.05 < P* < 0.1.

To test whether age-related oxidative stress directly disrupts NAD^⁺^ and adenine metabolism, young COCs were exposed to H₂O₂ for 30 min to induce oxidative stress prior to IVM in the presence or absence of NMN (Fig. 7a). H₂O₂ treatment had no detectable effect on NAD^⁺^ metabolites within MII oocytes (Fig. 7b). NMN supplementation significantly increased NMN levels in control oocytes, but also reduced NADPH, suggesting that exogenous NMN alters the balance of NAD⁺-related metabolites in MII oocytes even in the absence of oxidative stress. Under H₂O₂-induced oxidative stress, NMN treatment was associated with trends toward increased NMN and NAD⁺ abundance in MII oocytes. In contrast, matched cumulus cells were more metabolically responsive to oxidative stress. H₂O₂ reduced NAD⁺ and meNAM, with trends toward reduced NADP^⁺^ and an increased NADPH/NADP^⁺^ ratio in both untreated and NMN-treated groups (Fig. 7c). Similarly, H₂O₂ treatment had no detectable effect on adenine metabolites within MII oocytes (Fig. 7d); however, trends toward decreased AMP and ATP, and an increase in the ATP/AMP ratio, were apparent in cumulus cells (Fig. 7e). These oxidative stress-associated changes were restored by NMN treatment, with metabolites returning toward levels observed in NMN-treated control cumulus cells. In contrast, in NMN-treated H₂O₂-exposed cumulus cells, NADH was significantly reduced, the NAD⁺/NADH ratio showed a trend toward an increase, and the NADPH/NADP^⁺^ ratio remained elevated relative to NMN-treated controls. This suggests that oxidative stress continues to alter elements of cumulus cell redox metabolism despite NMN supplementation.

**Fig. 7:**
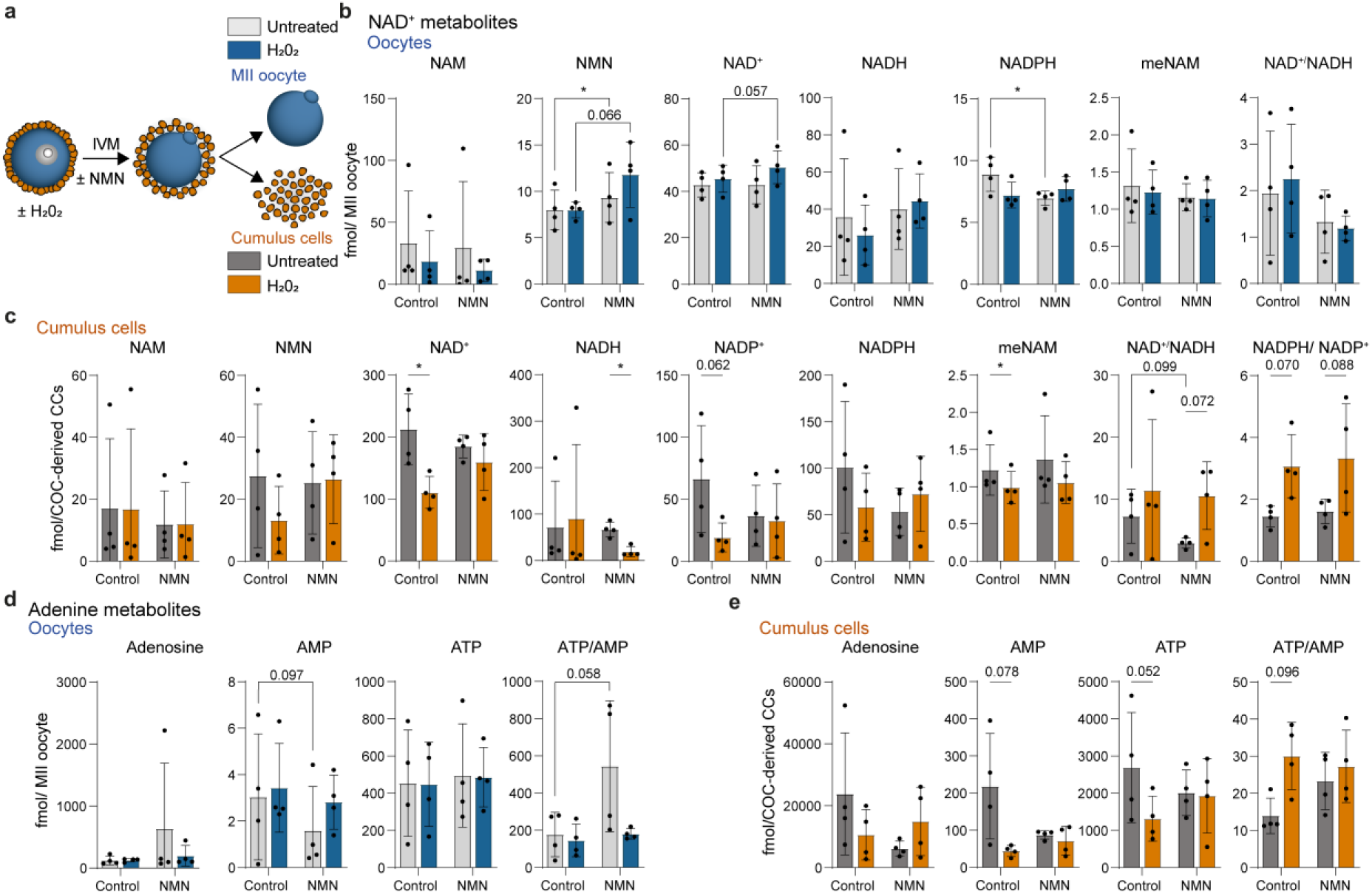
NMN reduces oxidative stress–induced disruption of NAD⁺ and adenine nucleotide metabolism in cumulus cells. **a,** Experimental overview. Germinal vesicle (GV)-stage cumulus–oocyte complexes (COCs) from reproductively young mice (5–7 wk) were treated with 100 µM hydrogen peroxide (H₂O₂) for 30 min before in vitro maturation (IVM) to the metaphase II (MII) stage in the presence or absence of nicotinamide mononucleotide (NMN). MII COCs were mechanically dissociated, and oocytes and matched cumulus cells were isolated for targeted LC–MS/MS. **b,c,** NAD⁺-related metabolites measured in **b,** MII oocytes and **c,** matched cumulus cells. **d,e,** Adenine metabolites measured in the same **d,** MII oocytes and **e,** matched cumulus cells. Metabolite abundance is shown as fmol per MII oocyte for oocyte samples and fmol per COC-derived cumulus cells (CCs) for cumulus cell samples. Bars show mean ± s.d.; individual dots represent independent biological replicates (n = 4). Each biological replicate comprised pooled oocytes or matched cumulus cells from 20–30 COCs collected from a minimum of 12 mice. Statistical significance was determined using repeated-measures two-way ANOVA, with H₂O₂ and NMN treatment as factors, biological replicate as the matching factor, and Geisser–Greenhouse correction applied, followed by Fisher’s LSD test for selected row- and column-wise pairwise comparisons. Asterisks indicate statistically significant selected comparisons: \**P* < 0.05. Exact P values are shown for selected comparisons with *0.05 < P < 0.1*.

NMN supplementation following oxidative insult partially rescued H₂O₂-induced reductions in polar body extrusion and improved redox homeostasis, reducing ROS accumulation and increasing glutathione-associated antioxidant capacity in cumulus-enclosed oocytes (Fig. 8a–d). NMN supplementation increased mitochondrial membrane potential in both untreated and H₂O₂-exposed oocytes (Fig. 8e), suggesting that NMN can enhance mitochondrial polarization in young COCs. H₂O₂ alone did not induce overt mitochondrial depolarization, distinguishing this acute oxidative stress model from the reproductive aging model, and may reflect the presence of relatively intact cumulus cell metabolic support in young H₂O₂-treated COCs.

**Fig. 8:**
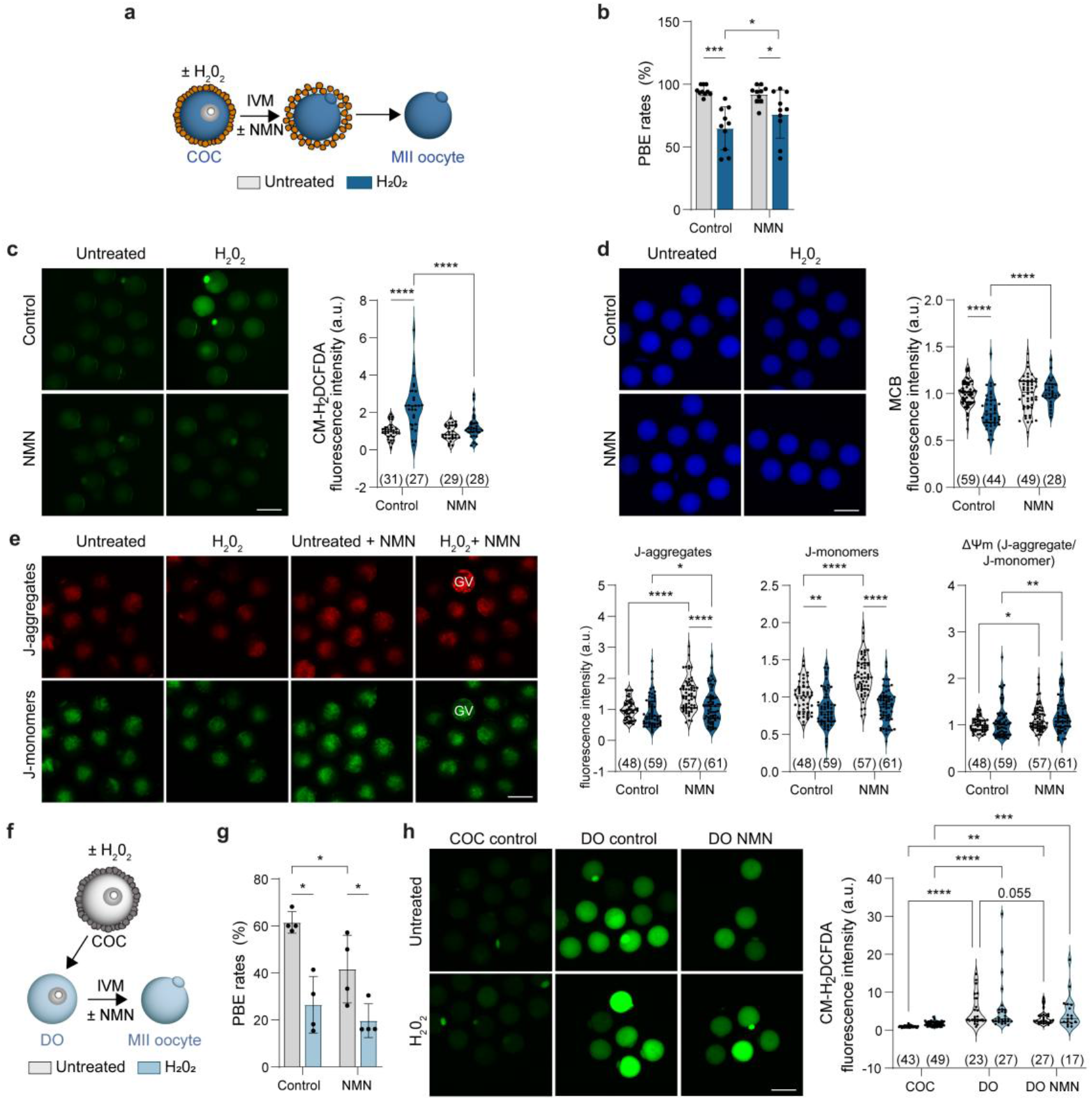
NMN mitigates acute oxidative stress in cumulus-enclosed oocytes. **a,** Experimental overview. Germinal vesicle (GV)-stage cumulus–oocyte complexes (COCs) from reproductively young mice (5–7 wk) were treated ± 100 µM hydrogen peroxide (H₂O₂) for 30 min before in vitro maturation (IVM) in the presence or absence of nicotinamide mononucleotide (NMN). **b,** Polar body extrusion (PBE) rates; **c,** CM-H₂DCFDA fluorescence intensity (indicative of reactive oxygen species); and **d,** monochlorobimane (MCB) fluorescence intensity (indicative of reduced glutathione) in metaphase II (MII) oocytes from COCs following oxidative stress and IVM with or without NMN treatment. **e,** JC-1 red J-aggregate fluorescence, green monomer fluorescence and red:green fluorescence ratio in MII oocytes from COCs following oxidative stress and IVM with or without NMN treatment. An arrested GV-stage oocyte visible in the representative JC-1 red J-aggregate fluorescence image is labeled “GV” for clarity and was excluded from quantification; only MII oocytes were included in quantitative analysis. **f,** GV-stage denuded oocytes (DOs) from reproductively young mice (5–7 wk) were treated ± 100 µM hydrogen peroxide (H₂O₂) for 30 min before in vitro maturation (IVM) in the presence or absence of nicotinamide mononucleotide (NMN). **g,** PBE rates; and **h,** CM-H₂DCFDA fluorescence intensity in MII-stage oocytes from denuded GV-stage oocytes, following oxidative stress and IVM with or without NMN treatment. Scale bars, 100 µm. Fluorescence intensity is shown in arbitrary units (a.u.) for all fluorescence analyses. Bars show mean ± s.d. For PBE analyses, each dot represents an independent biological replicate, with each replicate containing a minimum of 10 oocytes. For fluorescence analyses, each dot represents an individual oocyte from a minimum of three biological replicates, with each replicate containing a minimum of four imaged oocytes. Oocyte numbers are annotated on the x-axis within each graph. For all experiments, oocytes were collected from a minimum of three mice. Statistical significance was determined using ordinary two-way ANOVA for analyses in c–g, with H₂O₂ and NMN treatment as factors, followed by Fisher’s LSD test for selected row- and column-wise pairwise comparisons. For CM-H₂DCFDA fluorescence intensity analysis in h, statistical significance was determined using ordinary two-way ANOVA, with H₂O₂ and NMN treatment as factors, followed by Fisher’s LSD test for selected row- and column-wise pairwise comparisons. Asterisks indicate statistically significant selected comparisons: \**P* < 0.05, \*\**P* < 0.01, \*\*\**P* < 0.001 and \*\*\*\**P* < 0.0001. Exact *P* values are shown for selected comparisons with *0.05 < P* < 0.1.

Denuded oocytes displayed elevated ROS and reduced polar body extrusion following IVM compared with cumulus-enclosed oocytes, indicating that removal of cumulus cell support alone compromises oocyte redox balance and maturation competence (Fig. 8f–h). H₂O₂ exposure did not further increase ROS in denuded oocytes, suggesting that denudation induces a high-ROS state during maturation. In contrast to cumulus-enclosed oocytes, NMN did not rescue polar body extrusion in denuded oocytes, with polar body extrusion remaining similar to H₂O₂-treated denuded oocytes (Fig. 8g). However, NMN-treated DOs showed a trend toward reduced ROS relative to control DOs (Fig. 8h). This indicates that NMN-mediated reduction of elevated ROS during maturation depends, at least in part, on the presence of cumulus cells. NMN-treated denuded oocytes also displayed reduced polar body extrusion rates relative to untreated denuded oocytes. When considering the accumulation of unmetabolized NMN in oocytes lacking cumulus cell contact (Fig. 4c), this suggests that NMN which cannot be converted to NAD⁺ is not metabolically inert.

While the overall metabolic trends were broadly consistent across experiments, minor differences were observed between control and NMN-treated MII oocytes and cumulus cells between our mouse models; SwissTacAusb (Fig. 5) and C57BL/6JAusb (Fig. 7). When directly comparing MII oocytes and cumulus cells between mouse models, we observed minor differences, including reduced NAD⁺, trends toward increased NADPH and an increased NADPH/NADP⁺ ratio in C57BL/6JAusb MII oocytes, and an increased NADPH/NADP⁺ ratio in C57BL/6JAusb cumulus cells (Extended Data Fig. 4). These subtle differences may contribute to minor variances observed between aging and oxidative stress datasets.

## Discussion

Declining oocyte NAD⁺ has emerged as a key contributor to female reproductive aging, yet the cause of this decline within the oocyte remains unresolved. Oocyte aging is usually considered through the lens of the oocyte as an exceptionally long-lived autonomous unit. Here, we show that reproductive aging alters cumulus cell NAD⁺ metabolism in both mice and humans, and that this somatic compartment is required to maintain oocyte NAD⁺ homeostasis through a previously unrecognized mechanism of somatic–germline coupling. Further, we show that NAD^⁺^ metabolism-mediated resolution of the age-related increase in ROS in oocytes is cumulus cell dependent. Overall, these findings reveal impaired cumulus cell metabolic support as a mechanism contributing to oocyte NAD⁺ decline and reframe oocyte aging as a failure of somatic–germline metabolic cooperation, and not only an intrinsic deterioration of the long-lived oocyte.

Previous studies have described declining NAD⁺ abundance in the whole ovary^20,22,36^ and reduced NAD⁺ metabolites in mouse^18,19^ and human^21^ oocytes with reproductive age. Our findings advance this by demonstrating that reproductive aging alters NAD⁺ metabolism to a greater extent in the oocyte’s somatic compartment, the cumulus cells, shifting the focus from the oocyte alone to the COC as a metabolic unit.

A central finding of this study is that somatic-germline metabolic cooperativity is required for oocyte NAD⁺ synthesis, which in turn is required for the oocyte to combat oxidative stress. Cumulus–oocyte communication lies at the heart of this metabolic symbiotic relationship. Isotope tracing showed that d₄-NMN could still enter and accumulate in oocytes after gap junctions were blocked with CBX treatment or oocytes were denuded, but its conversion into labeled NAD⁺ was impaired. Thus, despite NMN availability, oocytes are unable to efficiently synthesize NAD⁺ when communication with cumulus cells is disrupted. In contrast, cumulus cells retained the capacity to efficiently synthesize NAD⁺ from NMN, maintaining NAD⁺ levels after CBX treatment. This compartmental difference supports a model in which oocytes and cumulus cells function as a cooperative metabolic unit, but with distinct roles in NAD⁺ metabolism (Fig. 4). Moreover, this cooperation is disrupted with age, whereby the age-related decline in cumulus cell NAD⁺ metabolism demonstrated here, together with known age-related reductions in TZPs and gap-junction coupling^40^, appear to limit the capacity of cumulus cells to metabolically support the oocyte. This model of intercellular metabolic cooperativity also resembles a similar observation in neurons, where NMN can be donated from damaged neurons to neighboring healthy neurons, which convert this NMN into NAD^⁺^ and export it back to the damaged neuron – thereby evading the NMN-sensitive SARM1 death pathway^58^. Notably, this bidirectional transport of metabolites occurs *via* gap junctions. Our data are consistent with a model whereby NMN can enter oocytes, but undergoes retrograde transport back into cumulus cells for its incorporation into NAD^⁺^, which is then transported back into the oocyte as required.

We and others have reported that supplementation of NAD⁺ precursors increased ovarian NAD⁺ abundance and reduced features of ovarian aging^18,20,22^. A striking feature of this study is that the metabolic benefit of exogenous NMN was most clearly reflected in the oocyte, even when cumulus cell NAD⁺ metabolism was not broadly restored. With reproductive aging, where oocyte NAD⁺ abundance and redox outcomes were compromised, NMN improved oocyte NAD⁺ abundance and homeostasis but did not broadly restore cumulus cell NAD⁺ metabolites. In contrast, during acute H₂O₂-induced oxidative stress, where the oocyte metabolome was relatively preserved, NMN partially restored cumulus cell metabolic changes as well as oocyte redox outcomes. These findings raise the intriguing possibility that metabolic support within the COC is directed toward preserving oocyte NAD^⁺^ homeostasis.

This notion of germ-cell metabolic sparing appears further exacerbated by meiotic maturation in aged COCs. In young oocytes, meiotic maturation was associated with reduced NAD⁺ metabolites, consistent with dynamic remodeling of the oocyte NAD⁺ pool during meiosis. These maturation-associated changes were less evident in aged oocytes; however, cumulus cell NAD⁺ and NADP⁺ levels were already compromised by aging in immature COCs, and by the end of oocyte meiotic maturation there was broad depletion of the NAD⁺ metabolites in cumulus cells from old mice. That pattern is consistent with the possibility that the cumulus cell NAD^⁺^ metabolome is depleted during maturation in favor of preserving NAD^⁺^ homeostasis in the germ cell.

Not only does the transfer of metabolites rely on cumulus cells, we further demonstrated that the redox benefit of NMN depends on cumulus cells. Denuded oocytes displayed elevated ROS and reduced polar body extrusion after IVM relative to cumulus-enclosed oocytes, demonstrating that removal of cumulus cell support alone causes oxidative stress. In contrast to cumulus-enclosed oocytes, NMN did not significantly reduce ROS or increase polar body extrusion in denuded oocytes. Thus, NMN-mediated resolution of elevated ROS during oocyte maturation requires, at least in part, an intact cumulus cell compartment. This cumulus cell dependence may reflect reduced access to cumulus-derived NAD⁺ metabolic support through NADP⁺/NADPH. As NADPH provides the reducing power required to regenerate reduced glutathione^25,26^, disrupted access during aging, where NAD⁺ metabolites are decreased or where cumulus–oocyte communication is lost, may limit the capacity of aged oocytes to maintain GSH-dependent antioxidant defense and protect against ROS.

These findings also have implications for IVM and therapeutic strategies aimed at improving oocyte quality with age. IVM is a non-experimental, but niche, infertility treatment and older patients in particular would benefit from an improvement in its efficacy^59^. NAD⁺ precursor supplementation has been explored as an approach to improve oocyte quality during IVM^60–63^, including in rescue-IVM, where immature oocytes collected after controlled ovarian stimulation have failed to mature by retrieval and are stripped of cumulus cells before culture. In this context, NMN treatment has been reported to modestly increase maturation rates in oocytes from women of advanced maternal age^63^. However, our findings support the efficacy of precursor supplementation of NAD⁺ metabolism to whole COCs during IVM as a potential target for improving the age-related decline in oocyte quality. Importantly, they demonstrate that NAD⁺ precursor supplementation should be considered in the context of cumulus–oocyte metabolic cooperation.

It remains unclear why oocytes have a limited capacity to convert NMN into NAD^⁺.^ One possibility is that adenine nucleotide metabolism represents a component of cumulus-cell support required for oocyte NAD⁺ synthesis. NAD⁺ biosynthesis depends not only on NMN availability but also on ATP, because nicotinamide mononucleotide adenylyltransferase (NMNAT) enzymes catalyze the ATP-dependent conversion of NMN to NAD⁺^48^. In our study, disruption of cumulus–oocyte communication in young COCs reduced AMP in both oocytes and cumulus cells, while ATP abundance was preserved. As ATP, adenosine diphosphate (ADP), and adenosine monophosphate (AMP) are regulated as an interconnected adenylate pool through adenylate kinase-mediated cycling^50,64^, the reduction in AMP despite maintained levels of ATP may reflect altered adenine nucleotide turnover rather than depletion of the total ATP pool. This altered adenylate balance may, in turn, constrain ATP-dependent NMNAT-mediated conversion of NMN to NAD⁺^48^.

This interpretation is consistent with previous work from our team, which showed that cumulus–oocyte communication supports adenine nucleotide flux into the oocyte. Exogenous ¹³C₅-AMP was taken up more efficiently by cumulus-enclosed oocytes than denuded oocytes, generating higher ¹³C₅-ADP and ¹³C₅-ATP, while gap-junction inhibition reduced ¹³C₅-ATP generation from ¹³C₅-AMP^65^. Notably, increasing ¹³C₅-AMP availability altered the proportion of labeled ATP relative to endogenous ATP, but did not change total oocyte ATP abundance^65^. This indicates that bulk ATP abundance can remain stable despite substantial changes in adenine nucleotide flux. However, oocytes matured in vitro without cumulus cells become ATP-depleted^66^, indicating that cumulus cell support does become limiting under the high ATP demand of maturation.

In the context of reproductive aging, oocyte adenine nucleotide changes were modest, with AMP altered only when maturation stage was considered (Extended Data Fig. 1). Reduced AMP has been similarly reported in human oocytes with increasing reproductive age^21^. In contrast, adenosine abundance was reduced in aged mouse and human cumulus cells, while ATP was also reduced in aged mouse cumulus cells. These changes may limit the capacity of cumulus cells to support adenine nucleotide homeostasis in the oocyte, with potential consequences for ATP-dependent NAD⁺ biosynthesis. However, it is important to note that intracellular ATP measurements are sensitive to sample collection and processing, with previous studies of human cumulus cells reporting decreased^67,68^, unchanged^69^ or increased^21^ ATP abundance with reproductive age. Together, these findings suggest that disruption of adenine nucleotide turnover contributes to impaired oocyte NAD⁺ homeostasis during reproductive aging.

In our working model, reduced NAD⁺ synthesis in aged oocytes and following disruption of cumulus–oocyte communication may reflect diminished cumulus-cell support for both adenine nucleotide balance and NAD⁺ precursor metabolism. During aging, where there is already a loss of cumulus–oocyte communication^40^, age-associated changes in cumulus-cell adenine nucleotide metabolism may further reduce the capacity for the oocyte to sustain ATP-dependent NMNAT activity, while altered NAD⁺ metabolism within aged cumulus cells may limit the generation or transfer of metabolites required for oocyte NAD⁺ homeostasis^33,34,58^. During reproductive aging, these defects in metabolic coupling converge to impair NAD⁺ synthesis in aged oocytes. An alternate possibility for future investigation is that NMNAT enzymes are more highly expressed in cumulus cells relative to oocytes, so that both the upper and lower bounds of oocyte NAD^⁺^ homeostasis can be regulated by the surrounding somatic cell compartment.

This study also raises several important new questions. Across experiments, the NAD^⁺^ metabolome of young oocytes and cumulus cells was insensitive to NMN supplementation, suggesting that NAD⁺ levels are tightly regulated. Indeed, the labeling experiments (Fig. 4) demonstrated that exogenous NMN entered or accumulated in oocytes without proportional increases across the broader NAD⁺ metabolite pool, raising the possibility that excess metabolites are diverted into terminal products such as meNAM, or released into the culture medium. Defining how the cumulus–oocyte complex buffers NAD⁺ levels would be an interesting avenue for future work. In addition, while this study identifies disrupted adenine/NAD⁺ metabolism as one limiting factor for oocyte NAD⁺ synthesis, broader classes of cumulus-derived signals undoubtedly also contribute to oocyte aging. In addition to metabolites, there is a growing body of literature identifying the intercellular transfer of larger cargo, within the cumulus–oocyte complex, including mRNA and long non-coding RNA^70,71^ and lipids^72^, an area that is being actively explored by us and others. It is possible that additional factors beyond the metabolites identified here are also transferred between cumulus cells and the oocyte, and that age-related changes in these factors will soon emerge as contributors to oocyte aging.

It is also important to acknowledge the study limitations. For human analyses, we were restricted to cumulus cells, as the number of matched human oocytes required for metabolomics was not feasible. Natural variability within human samples was dictated by individual patient health, disease status and treatment modality. In addition, low oocyte yield, particularly from reproductively aged animals, limited sample size and meant that not all metabolites were consistently detectable across all experiments. We also did not assess metabolites released into the culture medium to determine the fate of excess NAD⁺ precursors. Finally, while the overall metabolic trends were broadly consistent across models, subtle model-dependent differences should be considered when comparing the aging and oxidative stress datasets. Nevertheless, ultra-sensitive targeted mass spectrometry generated biologically meaningful patterns across mouse and human samples.

Taken together, our study reframes the age-related decline in oocyte NAD⁺ not as a purely cell-autonomous defect, but rather as a failure of somatic–germline metabolic coupling. We identify declining NAD⁺ metabolism in cumulus cells, demonstrate that cumulus cells are critical regulators of oocyte NAD⁺ homeostasis, and show that this somatic compartment is required for NAD⁺ metabolism to support redox balance during oocyte maturation. Importantly, these findings position cumulus cells not simply as passive support cells, but as an active metabolic unit that is metabolically coupled to the oocyte. These results provide new insight into the metabolic regulation of oocyte aging and highlight this new model of shared intercellular NAD⁺ metabolism within the cumulus–oocyte complex during IVM as a potential target for improving assisted reproductive outcomes.

## Materials and Methods

### Animals

All animal experiments were approved by the University of New South Wales Animal Ethics Committee (UNSW AEC; 23/25A, 22/40A, iRECS 8829, and iRECS 6531) and were conducted in accordance with the Australian Code for the Care and Use of Animals for Scientific Purposes. Female C57BL/6JAusb mice and SwissTacAusb mice were purchased from Australian Bio-Resources (ABR; Moss Vale, NSW, Australia). Animals were allowed to acclimatize for 1 week before use. Ex-breeder SwissTacAusb mice were housed for up to 6 months in the UNSW Lowy Animal Facility until they reached 12–14 months of age. Mice were maintained in individually ventilated cages at 22°C and 50–70% humidity at a density of up to five mice per cage, under a 12 h light/dark cycle, with *ad libitum* access to food and water. Mice were humanely killed *via* cervical dislocation.

### Human cumulus cell collection

Approval of this study was granted by the South Eastern Sydney Local Health District Ethics Committee (2023/ETH01654). Cumulus cells discarded during routine ICSI were donated by patients undergoing treatment at the Fertility & Research Centre (FRC) in the Royal Hospital for Women, Sydney. Informed consent for the use of cumulus cells for research was obtained. Ovarian stimulation was performed with FSH using either gonadotropin-releasing hormone (GnRH) agonist or antagonist protocols. Following collection of COCs, cumulus cells and oocytes were separated by incubation with SynVitro Hyadase (15115001, ORIGIO, Måløv, Denmark) in Oil for Tissue Culture (ART-4008-P, SAGE In-Vitro Fertilization, Inc., Trumbull, CT, USA) and were further mechanically separated by pipetting through handling media (83100125, ORIGIO) until the oocytes were fully isolated. Cumulus cells were maintained at 37°C and collected approximately 1–2 h after isolation. Cells were transported at 37°C in a portable incubator for approximately 15 min to the research laboratory. Cumulus cells from 23 patients undergoing routine ICSI treatment between November 2025 and February 2026 were included in this study. Patients undergoing fertility preservation for medical indications, including cancer treatment, or with known diagnoses of polyendocrine metabolic ovarian syndrome (PMOS), premature ovarian insufficiency (POI) or other systemic illness, were excluded from the analysis. Only cumulus cell samples containing >5 COCs, >100,000 cells, with >50% viability and no visible blood contamination, were included for analysis; all included samples were processed using a uniform extraction protocol.

### Mouse oocyte and cumulus–oocyte complex collection

To collect immature germinal vesicle (GV)-stage oocytes, COCs were collected from ovaries of 5–7-week-old C57BL/6JAusb females. Mice were administered an intraperitoneal injection of 5 IU pregnant mare serum gonadotropin (PMSG; Prospec-Tany Technogene Ltd, Israel). Mice were humanely killed 46–48 h after PMSG injection, ovaries were collected, and COCs were released into HEPES-buffered handling medium. Handling medium was prepared using custom α-minimum essential medium powder (α-MEM; 8.8 g/L; ME14196P1, Gibco, Thermo Fisher Scientific, Waltham, MA, USA), formulated with 2 mM L-alanyl-L-glutamine in place of L-glutamine, and supplemented with 6 mM sodium bicarbonate (S8875, Sigma-Aldrich, St Louis, MO, USA), 10 mM HEPES free acid (H6147, Sigma-Aldrich), 10 mM HEPES sodium salt (H7006, Sigma-Aldrich), 50 µg/mL gentamicin (G1914, Sigma-Aldrich) and 3 mg/mL bovine serum albumin (BSA; A7906, Sigma-Aldrich). Handling medium was further supplemented with 100 µM 3-isobutyl-1-methylxanthine (IBMX; I5879, Sigma-Aldrich) to prevent oocyte maturation. COCs were released from preovulatory follicles using a 29-gauge needle and collected using flame-pulled borosilicate Pasteur pipettes. COCs were selected based on the presence of a fully enclosed oocyte surrounded by intact, compact cumulus cells. For experiments involving aged animals, neither young nor old mice were hormonally stimulated prior to collection. Depending on the experiment, GV COCs were either processed immediately, or subjected to culture with or without meiotic arrest.

### Oocyte and COC culture

Isolated COCs containing GV oocytes (intact), and isolated GV oocytes (denuded) were cultured in culture medium for 17 h at 37°C in humidified air containing 5% CO₂. Culture medium was prepared using custom α-minimum essential medium powder (α-MEM; 8.8 g/L; ME14196P1, Gibco, Thermo Fisher Scientific), formulated with 2 mM L-alanyl-L-glutamine in place of L-glutamine, and supplemented with 26 mM sodium bicarbonate (S8875, Sigma-Aldrich), 50 µg/mL gentamicin (G1914, Sigma-Aldrich), 3 mg/mL fatty acid-free BSA (AlbumiNZ; 02199899, MP Biomedicals, New Zealand) and 1 mg/mL fetuin (F3385, Sigma-Aldrich). Where indicated, culture medium was further supplemented with 100 µM IBMX. Gap junction communication within COCs was disrupted using 200 µM carbenoxolone (CBX; J63714.06, Thermo Fisher Scientific). Intact, denuded, and CBX-disrupted COCs were supplemented with or without 500 µM NMN (GeneHarbor Biotechnology, Hong Kong, China). For isotope-tracing experiments, 500 µM nicotinamide-d₄ mononucleotide (NMN-d₄, TRC-N407768; Toronto Research Chemicals) was used. After 17 h, COCs were mechanically separated into oocytes and cumulus cells using flame-pulled borosilicate Pasteur pipettes prior to downstream analysis.

### In vitro maturation

COCs were washed five times in 3 mL of IBMX-free culture medium to ensure complete removal of IBMX prior to *in vitro* maturation (IVM) and cultured for 17 h in culture media supplemented with 50 ng/mL mouse amphiregulin (mAREG; 989-AR, R&D Systems), and 50 ng/mL mouse epiregulin (mEREG; 1068-ER, R&D Systems) in 4-well culture dishes (NUNC, 179830, Thermo Fisher Scientific) at 37°C, 5% CO₂ in humidified air. During IVM, COCs were incubated with or without 500 µM NMN or NMN-d₄. After 17 h, expanded COCs were removed from culture and MII oocytes and cumulus cells were mechanically separated. Oocytes were morphologically assessed as GV by intact nuclear envelope and as MII by polar body extrusion.

### Induction of oxidative stress

COCs were treated with 100 µM hydrogen peroxide (H₂O₂; AJA260-500, Sigma-Aldrich) for 30 min at 37°C in handling medium supplemented with 100 µM IBMX under paraffin oil as previously described^55,73^. Following H₂O₂ treatment, COCs underwent IVM as described above.

### Assessment of intracellular reactive oxygen species, glutathione, and mitochondrial membrane potential

Following IVM, expanded COCs were mechanically denuded, and oocytes were used for assessment of intracellular reactive oxygen species (ROS), glutathione (GSH) levels, and mitochondrial membrane potential. For ROS detection, oocytes were incubated in 4 µM chloromethyl-2′,7′-dichlorodihydrofluorescein diacetate (CM-H2DCFDA; C6827, Thermo Fisher Scientific) for 1 h as previously described^55^. For GSH assessment, oocytes were incubated in 12.5 µM monochlorobimane (MCB; M1381MP, Thermo Fisher Scientific) for 30 min. For mitochondrial membrane potential, oocytes were incubated in JC-1 dye (T3168, Invitrogen, Thermo Fisher Scientific, Waltham, MA, USA) at 1 µg/mL for 30 min. All incubations were performed in HEPES-buffered α-MEM supplemented with 3 mg/mL BSA at 37°C, protected from light under paraffin oil. Following staining, oocytes were washed through three 100 µL droplets of fresh medium to remove excess dye and transferred into 5 µL droplets of medium overlaid with paraffin oil on glass-bottom imaging dishes (FD35-100, World Precision Instruments, Sarasota, FL, USA). MCB–GSH adduct fluorescence was detected using a DAPI filter set (Ex/Em: 358/461 nm) on an EVOS FL fluorescence microscope (Thermo Fisher Scientific, Waltham, MA, USA). ROS-dependent oxidation of CM-H₂DCFDA to DCF was detected at Ex/Em: 488/520 nm using either an EVOS FL fluorescence microscope or a ZEISS LSM 990 confocal microscope (Carl Zeiss Microscopy, Jena, Germany). JC-1 fluorescence was detected in both the green monomer channel (Ex/Em: 514/529 nm) and red J-aggregate channel (Ex/Em: 514/590 nm) using a ZEISS LSM 900 or ZEISS LSM 990 confocal microscope. For confocal imaging, whole oocytes were captured as z-stacks with a 2.5 µm step size. Mitochondrial membrane potential was expressed as the ratio of red to green fluorescence intensity, with a decreased red:green ratio indicating mitochondrial depolarization. Relative fluorescence intensities of MCB–GSH, DCF, and JC-1 were quantified in individual oocytes using Fiji/ImageJ software (National Institutes of Health, USA). For z-stacks, sum-intensity projections were generated for quantification. Fluorescence intensity was calculated as corrected total cell fluorescence (CTCF), defined as: CTCF = integrated fluorescence intensity − (area of selected cell × average background fluorescence). For each biological replicate, a minimum of four individual oocytes per group were imaged and analyzed. Data from experimental replicates were normalized to appropriate untreated controls. All imaging and analysis were performed using identical acquisition settings and exposure times and applied consistently across experimental groups.

### LC–MS/MS sample preparation and nucleotide extraction

NAD⁺ and adenine metabolites were quantified using liquid chromatography with tandem mass spectrometry detection (LC–MS/MS), as previously described^74,75^ with minor modifications. To prepare samples, following collection and/or culture, denuded oocytes or intact COCs were washed five times in 3 mL of Dulbecco’s phosphate-buffered saline (DPBS; 14190144, Gibco, Thermo Fisher Scientific) supplemented with 0.2% polyvinyl alcohol (PVA; 360627, Sigma-Aldrich), hereafter referred to as PBS/PVA, at 37°C. To collect isolated cumulus cells derived from COCs, cumulus cells were mechanically denuded from COCs in ∼10 µL PBS/PVA in a 1.5 mL tube using a flame-pulled borosilicate Pasteur pipette. Oocytes were then removed from the tube. Any remaining cumulus cells were washed off the oocytes in 3 mL PBS/PVA, and oocytes were transferred into a 1.5 mL tube containing ∼10 µL PBS/PVA. All oocyte and cumulus cell samples were paired for metabolomics analysis. Human cumulus cells were collected directly from the FRC and washed three times in 1 mL of PBS/PVA, at 37°C, with centrifugation at 750 × g for 5 min at 37°C. Due to variability in human cumulus cell recovery per COC, cumulus cell numbers were counted using an automated cell counter (Countess 3, Thermo Fisher Scientific) for normalization during data analysis.

For nucleotide extraction for LC–MS/MS, 250 µL of ice-cold 80% methanol (LC-MS grade) was added directly to samples immediately after collection. Samples were sonicated on ice for 1 min (40% power, 45% pulsed) using an ultrasonic homogenizer to ensure complete cell lysis. Extracts were incubated at −80°C for 5–10 min to enhance protein precipitation, followed by centrifugation at 14,000 × g for 10 min at 4°C to pellet protein and debris. The supernatant was diluted to 40% methanol with ultrapure H₂O, mixed, and filtered through 3 kDa centrifugal filter units (Amicon Ultra, UFC500396, MilliporeSigma, Burlington, MA, USA) at 14,000 × g for 2.5 h at 4°C. The resulting filtrate was collected and stored at −80°C until the day of LC–MS/MS analysis. The sample eluate was evaporated using a Savant SpeedVac vacuum centrifuge for 2.5 h at the low heat setting, with the chamber temperature maintained at room temperature. Samples were resuspended in 100 mM ammonium acetate supplemented with a fixed volume of isotope-labeled internal standard mixture. The internal standard stock mixture contained approximately 20 µM each of ¹³C₅-adenosine, ¹³C₅-AMP, ¹⁵N₅-ADP, ¹³C₁₀¹⁵N₅-ATP, NAM-d₄, NMN-d₄, NAD-d₄ and NADH-d₅ (Toronto Research Chemicals, Canada). Samples were then filtered through a 0.22 µm filter.

For isotope-tracing experiments, in which NMN-d₄ was supplemented during culture, ¹⁵N₅-ADP was used as a non-overlapping stable isotope-labeled internal standard to avoid interference between deuterated internal standards and tracer-derived isotopologues.

### LC–MS/MS

Metabolites were measured using either a TSQ Altis Plus triple quadrupole mass spectrometer or a TSQ Vantage triple quadrupole mass spectrometer coupled to a Vanquish UPLC system (Thermo Fisher Scientific, Waltham, MA, USA), as previously described^74,75^ with minor modifications. Metabolites were separated on a NH₂ column (150 × 2.0 mm, 3 µm; 0F-4377-B0, Phenomenex, Torrance, CA, USA) using a binary solvent gradient consisting of 5 mM ammonium acetate (pH 9.5 adjusted with ammonium hydroxide; mobile phase A) and acetonitrile (mobile phase B), with a flow rate of 250 µL/min. The percentage of mobile phase B was set at 75% (0–1 min), 40% (2–6 min), 30% (7 min), 5% (8–18 min), and 75% (19–30 min). The injection volume was 20–25 µL, with a Q1 resolution of 0.7 and Q3 resolution of 1.2. Collision-induced dissociation gas (mTorr) was set at 1.5. Spectra were processed and peak areas integrated using Xcalibur software with Quan Browser (Thermo Fisher Scientific). The targeted metabolite panel comprised NAM, NMN, NAD⁺, NADH, NADP⁺, NADPH, methyl-nicotinamide (meNAM), adenosine, AMP, ADP and ATP. All metabolites were assessed across experiments. Metabolites that were below the limit of detection in the majority of samples across all groups within a given experiment were not included in downstream analysis for that experiment. For metabolites included in an analysis, individual samples with values below the limit of detection were imputed as the lowest detectable value for that metabolite within the corresponding experiment divided by √2 before normalization and statistical analysis. Optimal selected reaction monitoring (SRM) transitions were established before sample analysis using authentic standards infused into the LC flow via a T-junction. The selected transitions, collision energies and source RF/S-lens voltages are as previously described^74,75^.

Calibration curves of individual metabolites were constructed using peak area ratios (peak area of the metabolite divided by the peak area of the selected internal standard) of each calibrator versus its concentration. Calibration curves covered a concentration range of 1–1,000 nM. Metabolite concentrations in oocyte and cumulus cell extracts were obtained from these calibration curves. Data were normalized to oocyte, COC, or cumulus cell number. Cellular energy charge was calculated using the following equation: Energy charge = ([ATP] + 1/2[ADP]) / ([ATP] + [ADP] + [AMP])^64^.

### Statistics

All statistical analyses and graphs were generated using GraphPad Prism 10 software (San Diego). Normality was tested using the Shapiro–Wilk normality test. Data were analyzed using two-tailed t-tests or two-way ANOVAs. Paired tests were used where appropriate. Specific statistical tests used for each experiment are noted in the figure legends. Graphs were assembled using Adobe Illustrator.

## Data availability

All data reported in this paper will be shared by the lead contacts upon request. Any additional information required to reanalyze the data reported in this paper is available from the lead contacts upon request.

## Acknowledgements

We are especially grateful to the women who donated cumulus cells, and to the embryologists, including Oksana Struikhina, Liza Tilia and Sarah du Toit-Thompson; research nurses, including Pru Sweeten; and medical consultants at the Fertility & Research Centre, including William Ledger and Rebecca Deans. We thank Fabrizzio Horta for helpful discussions. We gratefully acknowledge the Bioanalytical Mass Spectrometry Facility (BMSF) for mass spectrometry support and the Katharina Gaus Light Microscopy Facility (KGLMF) for microscopy support.

## Author information

### Contributions

Conceptualization, B.P.M., L.E.W., and R.B.G.; methodology, B.P.M., S.B. and R.P.; formal analysis, B.P.M. and D.L.; investigation, B.P.M., D.L., E.R.F., A.V., D.W., and M.J.B.; resources, R.B.G. and B.P.M.; writing – original draft, B.P.M. with assistance from L.E.W. and R.B.G.; writing – review & editing, all authors; visualization, B.P.M., L.E.W., and R.B.G.; supervision, B.P.M., L.E.W., and R.B.G.; project administration, B.P.M., L.E.W., and R.B.G.; and funding acquisition, B.P.M. and R.B.G. All authors approved the final manuscript.

## Extended Data Figures and Figure Legends

**Extended Data Fig. 1:**
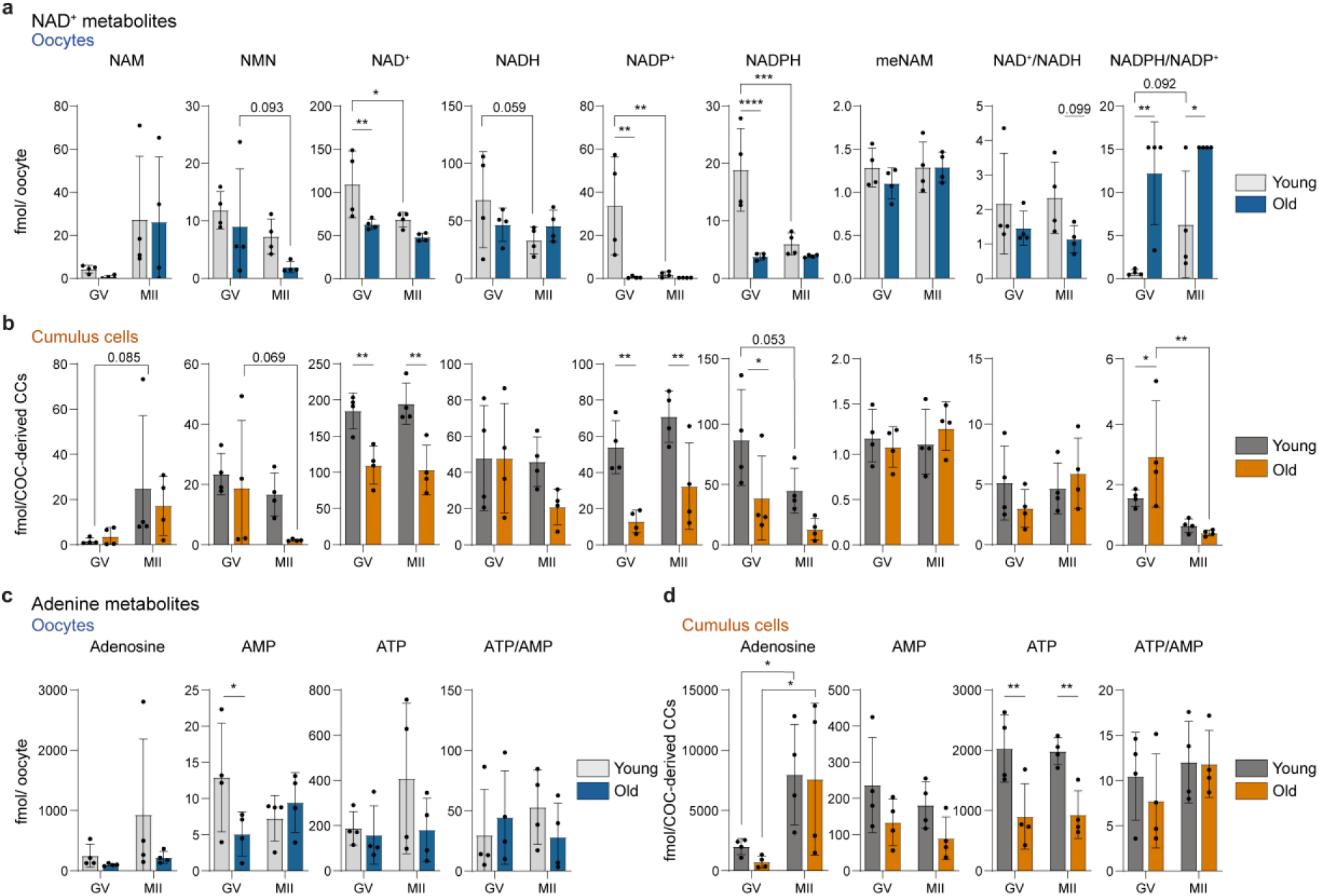
Reproductive aging alters NAD⁺ and adenine metabolite remodeling across oocyte meiotic maturation. Germinal vesicle (GV)-stage oocytes and matched cumulus cells were isolated from immature cumulus–oocyte complexes (COCs), and metaphase II (MII)-stage oocytes and matched cumulus cells were isolated from in vitro matured COCs from reproductively young (5–7 wk) and old (12–14 mo) mice. **a,b,** NAD⁺-related metabolites measured by targeted LC–MS/MS in **a,** oocytes and **b,** matched cumulus cells. **c,d,** Adenine metabolites measured by targeted LC–MS/MS in **c,** oocytes and **d,** matched cumulus cells. Metabolite abundance is shown as fmol per oocyte for oocyte samples and fmol per COC-derived cumulus cells (CCs) for cumulus cell samples. Bars show mean ± s.d.; individual dots represent independent biological replicates (n = 4). Each biological replicate comprised pooled oocytes or matched cumulus cells from 20–30 COCs collected from a minimum of three mice. GV- and MII-stage samples were collected as independent pooled biological replicates within the same experimental series, rather than paired samples from the same COCs; these datasets are also presented in Fig. 1 and Fig. 5, respectively. Statistical significance was determined using ordinary two-way ANOVA, with age and maturation stage as factors, followed by Fisher’s LSD test for selected row- and column-wise pairwise comparisons. Asterisks indicate statistically significant selected comparisons: \**P* < 0.05, \*\**P* < 0.01, \*\*\**P* < 0.001 and \*\*\*\**P* < 0.0001. Exact *P* values are shown for selected comparisons with *0.05 < P* < 0.1.

**Extended Data Fig. 2:**
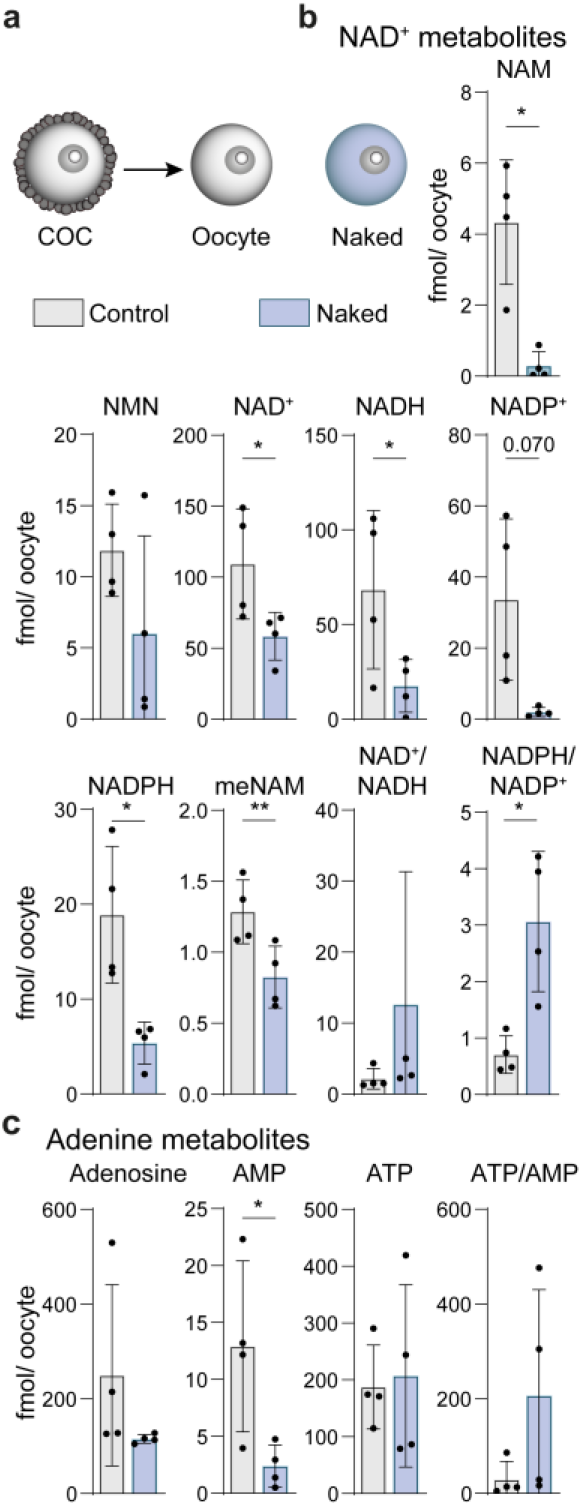
NAD⁺ and adenine metabolites differ between cumulus-enclosed and naked oocytes. **a,** Experimental overview. Germinal vesicle (GV)-stage oocytes from reproductively young mice (5–7 wk) were isolated from cumulus–oocyte complexes (COCs; control) and compared with oocytes collected naturally lacking surrounding cumulus cells (naked oocytes). **b,** NAD⁺-related metabolites measured in control and naked oocytes by targeted LC–MS/MS. **c,** Adenine metabolites measured in control and naked oocytes by targeted LC–MS/MS. Metabolite abundance is shown as fmol per oocyte. Bars show mean ± s.d.; individual dots represent independent biological replicates (n = 4). Each biological replicate comprised pooled oocytes from 20–30 COCs collected from a minimum of three mice. Comparisons were made between cumulus-enclosed control oocytes and naked oocytes within matched biological replicates. For normally distributed data, statistical significance was determined using two-sided paired t-tests. Where one or both groups failed normality testing, two-sided Wilcoxon matched-pairs signed-rank tests were used. Asterisks indicate statistical significance: \**P* < 0.05 and \*\**P* < 0.01. Exact P values are shown for comparisons with *0.05 < P* < 0.1; unlabeled comparisons had *P* ≥ 0.1.

**Extended Data Fig. 3:**
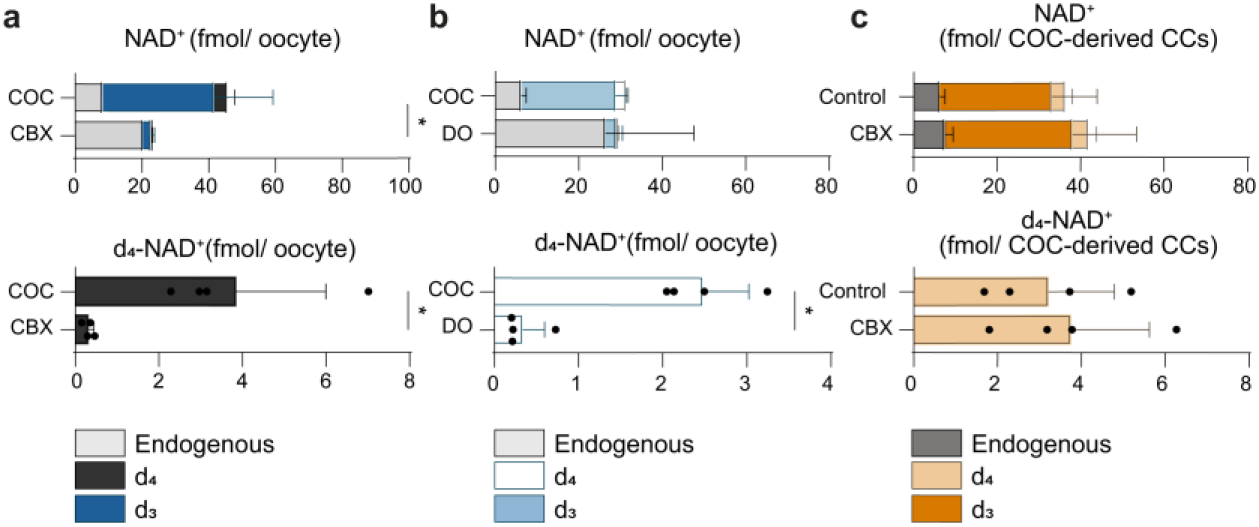
Total endogenous and deuterium-labeled NAD⁺ metabolite abundance following d₄-NMN tracing. Germinal vesicle (GV)-stage cumulus–oocyte complexes (COCs) from reproductively young mice (5–7 wk) were cultured intact as controls, or treated with carbenoxolone (CBX) to inhibit gap-junction communication, or mechanically denuded to remove cumulus cells (denuded oocytes; DOs), in the presence of deuterium-labeled NMN (d₄-NMN) for 17 h under isobutyl-1-methylxanthine (IBMX)-mediated meiotic arrest. **a,b,** Endogenous and deuterium-labeled NAD⁺ isotopologues (d₄ and d₃) measured by targeted LC–MS/MS in oocytes following **a,** CBX treatment or **b,** mechanical denudation. **c,** Endogenous and deuterium-labeled NAD⁺-related isotopologues (d₄ and d₃) measured by targeted LC–MS/MS in matched cumulus cells from COCs treated ± CBX. Metabolite abundance is shown as fmol per oocyte for oocyte samples and fmol per COC-derived cumulus cells (CCs). Bars show mean ± s.d.; individual dots represent independent biological replicates (n = 4). Each biological replicate comprised pooled oocytes or matched cumulus cells from 40–50 COCs collected from a minimum of six mice. For normally distributed data, statistical significance was determined using two-sided unpaired Welch’s t-tests. Where one or both groups failed normality testing, two-sided Mann–Whitney tests were used. Asterisks indicate statistical significance: \**P* < 0.05. Exact *P* values are shown for comparisons with *0.05 < P* < 0.1; unlabeled comparisons had *P* ≥ 0.1.

**Extended Data Fig. 4:**
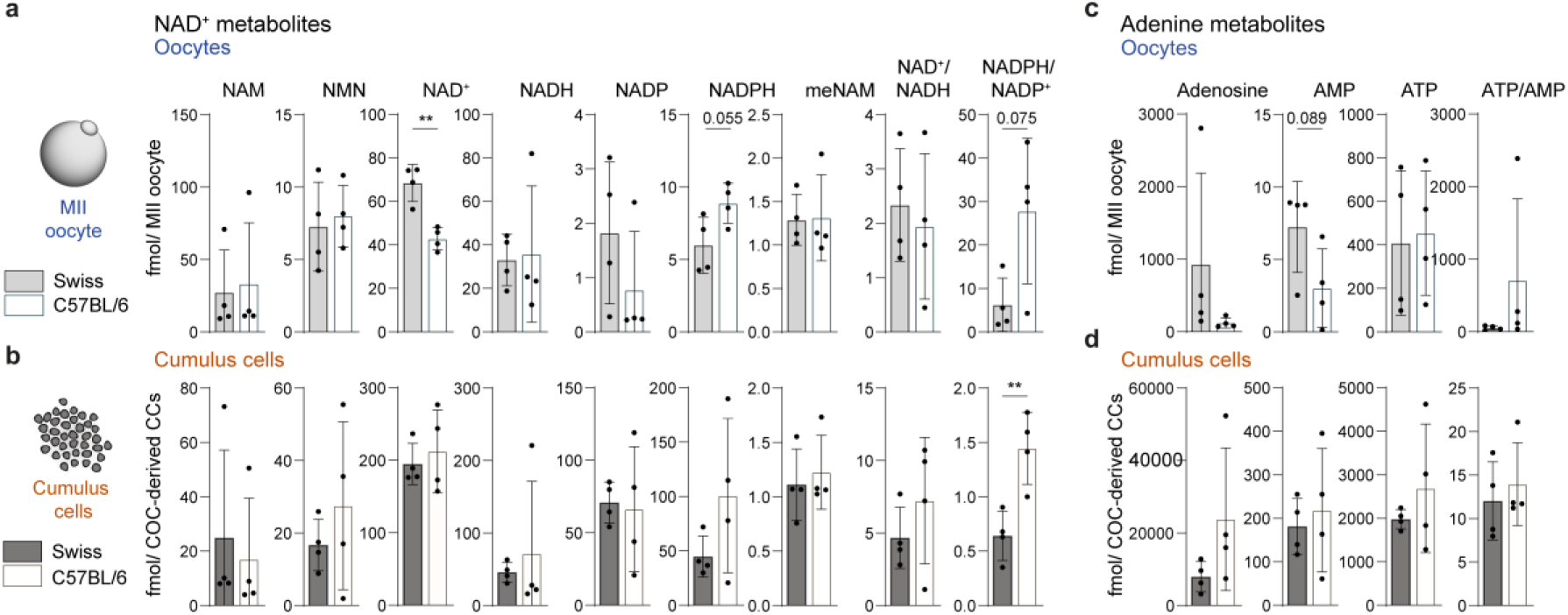
NAD⁺ and adenine metabolites in MII oocytes and cumulus cells from SwissTacAusb and C57BL/6JAusb mice. **a,b,** NAD⁺-related metabolites measured by targeted LC–MS/MS in **a,** metaphase II (MII)-stage oocytes and **b,** matched cumulus cells isolated from COCs from reproductively young (5–7 wk) SwissTacAusb (Swiss) and C57BL/6JAusb (C57BL/6) mice. **c,d,** Adenine metabolites measured by targeted LC–MS/MS in **c,** MII oocytes and **d,** matched cumulus cells from the same experimental groups. Metabolite abundance is shown as fmol per MII oocyte for oocyte samples and fmol per COC-derived cumulus cells (CCs) for cumulus cell samples. Bars show mean ± s.d.; individual dots represent independent biological replicates (n = 4). Each biological replicate comprised pooled oocytes or matched cumulus cells from 20–30 COCs collected from a minimum of three mice. For normally distributed data, statistical significance was determined using two-sided unpaired Welch’s t-tests. Where one or both groups failed normality testing, two-sided Mann–Whitney tests were used. Asterisks indicate statistical significance: \**P* < 0.05 and \*\**P* < 0.01. Exact *P* values are shown for comparisons with *0.05 < P* < 0.1; unlabeled comparisons had *P* ≥ 0.1.

